# Assessing Conservation of Alternative Splicing with Evolutionary Splicing Graphs

**DOI:** 10.1101/2020.11.14.382820

**Authors:** Diego Javier Zea, Sofya Laskina, Alexis Baudin, Hugues Richard, Elodie Laine

## Abstract

Understanding how protein function has evolved and diversified is of great importance for human genetics and medicine. Here, we tackle the problem of describing the whole transcript variability observed in several species by generalising the definition of splicing graph. We provide a practical solution to building parsimonious *evolutionary* splicing graphs where each node is a minimal transcript building block defined across species. We show a clear link between the functional relevance, tissue-regulation and conservation of AS events on a set of 50 genes. By scaling up to the whole human protein-coding genome, we identify a few thousands of genes where alternative splicing modulates the number and composition of pseudo-repeats. We have implemented our approach in ThorAxe, an efficient, versatile, and robust computational tool freely available at https://github.com/PhyloSofS-Team/thoraxe. The results are accessible and can be browsed interactively at http://www.lcqb.upmc.fr/ThorAxe.

## Introduction

Eukaryotes have evolved a transcriptional mechanism that can augment the protein repertoire without increasing genome size. A gene can be transcribed, spliced, and matured into several transcripts by choosing different initiation/termination sites or by selecting different exons (Graveley, 2001). Alternative splicing (AS) concerns almost all multi-exon genes in vertebrates (Wang et al., 2008). It can affect transcript maturation and post-transcriptional regulation, or result in protein isoforms (“proteoforms”) displaying different shapes (Birzele et al., 2008), interactions partners (Yang et al., 2016), and functions (Baralle and Giudice, 2017; Kelemen et al., 2013). AS misregulation is associated with the development of cancer, among other diseases (Climente-González et al., 2017; Scotti and Swanson, 2016; Lim et al., 2011; Ward and Cooper, 2010; Wang and Cooper, 2007). Moreover, the influence of natural AS variations between human populations on disease susceptibility is increasingly recognised (Park et al., 2018). Hence, understanding how AS contributes to protein function diversification is of utmost importance for human genetics and medicine.

In recent years, the advent of high-throughput sequencing technologies like RNA-Seq has made possible in-depth surveys of transcriptome complexity (Wang et al., 2008; Sultan et al., 2008). However, evaluating how many of the detected transcripts are translated and functional in the cell remains challenging (Wang et al., 2018; Kim et al., 2014). This difficulty has stimulated the development of integrative approaches combining gene annotations, RNA-Seq data and also data generated by other high-throughput techniques (Marti-Solano et al., 2020; de la Fuente et al., 2020; Louadi et al., 2020; Ait-hamlat et al., 2020; Agosto et al., 2019; Sterne-Weiler et al., 2018; Denti et al., 2018; Tapial et al., 2017; Tranchevent et al., 2017; Weatheritt et al., 2016; Ezkurdia et al., 2015; Rodriguez et al., 2013; Gonzàlez-Porta et al., 2013; De La Grange et al., 2010) toward a better characterisation of the AS landscape and phenotypic outcome. Recent studies underscore AS functional impact and contribution to protein diversity (Marti-Solano et al., 2020; Agosto et al., 2019).

Evolutionary conservation can arguably serve as a reliable indicator of function. Indeed, we expect splice variants selected over millions of years of evolution to comply with physical and environmental constraints and thus to play a functional role. The classical approach for assessing AS evolutionary conservation first identifies orthologous exons between species and then compares their inclusion/exclusion rates across cell/tissue types. A common practice for orthology detection is the BLAST (Altschul et al., 1990) “all-against-all” methodology (Nichio et al., 2017). More specialised protocols based on pairwise genomic sequence alignments (Mei et al., 2017; Herrero et al., 2016; Abascal et al., 2015; Modrek and Lee, 2003; Xing and Lee, 2005), or multiple alignments of genomic or protein sequences (Szalkowski, 2012; Christinat and Moret, 2012) have been proposed to detect orthology relationships between exons. Challenges associated with this task include correctly handling large indel events, finding plausible matches for highly divergent sequences, and resolving ambiguities arising from highly similar sequences (*e.g.*, resulting from in-gene duplication) or very short sequences. To compare the alternative usage of orthologous exons, compact representations of transcript variability, such as splicing graphs (Heber et al., 2002), can be employed. The nodes of a splicing graph represent the exons, and the edges denote exon junctions. Hence, a way to assess AS conservation between two genes would be to compare the environment of their orthologous exons in the two corresponding splicing graphs. However, until now, there exists no method combining these two layers of information.

Here we developed a novel and general method that revisits splicing graph representation to account for the whole transcript variability observed in a set of species. Specifically, we map all transcriptomic information coming from many species on an *evolutionary splicing graph*, where the nodes represent minimal transcript building blocks defined across species. Formally, each node, which we call a *spliced-exon* (or *s-exon*), is a set of aligned protein exonic sequences coming from different species. We recast the problem of exon orthology detection in the context of AS as that of determining the most parsimonious evolutionary splicing graph. We show that our strategy accurately resumes intra- and inter-species transcript variability. We compile a curated set of AS events involved in function diversification and show that they display evolutionary conserved tissue-regulation patterns. At the human protein-coding genome scale, we provide, for the first time, granular estimates of AS evolutionary conservation and significantly improve our knowledge on the amount of AS that is functionally relevant. AS is conserved across a wide range of evolutionary distances, is not limited to ancient events, and does not generate conserved alternative isoforms in all of the proteins. Interestingly, our analysis reveals evolutionary conserved AS-induced modulation of pseudo-repeats in about 10% of the proteins in human.

Our method is freely available in the form of a stand-alone package and python module at https://github.com/PhyloSofS-Team/thoraxe. It is fully automated and rapid. It takes less than 1 minute to treat one gene across one dozen species, on average. We also provide to the community a carefully curated set of 50 genes for which there is experimental evidence of AS-induced function diversification. All data are accessible and can be browsed interactively at http://www.lcqb.upmc.fr/ThorAxe.

## Results

### Evolution-informed model describes transcript variability

Our method decomposes a set of transcripts observed in several species into minimal building blocks, the s-exons, and constructs a graph where each transcript is encoded as a path (**Fig. 1A-B**). Graphs are convenient compact representations of transcript variability. Given a gene *G_i_*, and its annotated transcripts described as sorted lists of genomic intervals, a *splicing graph* (*SG*) can be defined as 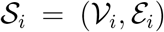. Each node 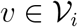 represents an exonic protein sequence occurring in at least one transcript isoform. As donor sites can be located inside an exon, the nodes of 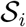 are not necessarily complete exons (**Fig. 1A**, see *n*_1_ and *n*_2_), and thus we call them *sub-exons*. The edge set 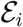 is inferred from the co-occurrences of sub-exons in the input transcripts. We distinguish the *structural* edges arising from intron boundaries (**Fig. 1A**, see *n*_1_ → *n*_3_) from the edges *induced* by the sub-exon boundaries (**Fig. 1A**, see *n*_1_ → *n*_2_).

**Figure 1:**
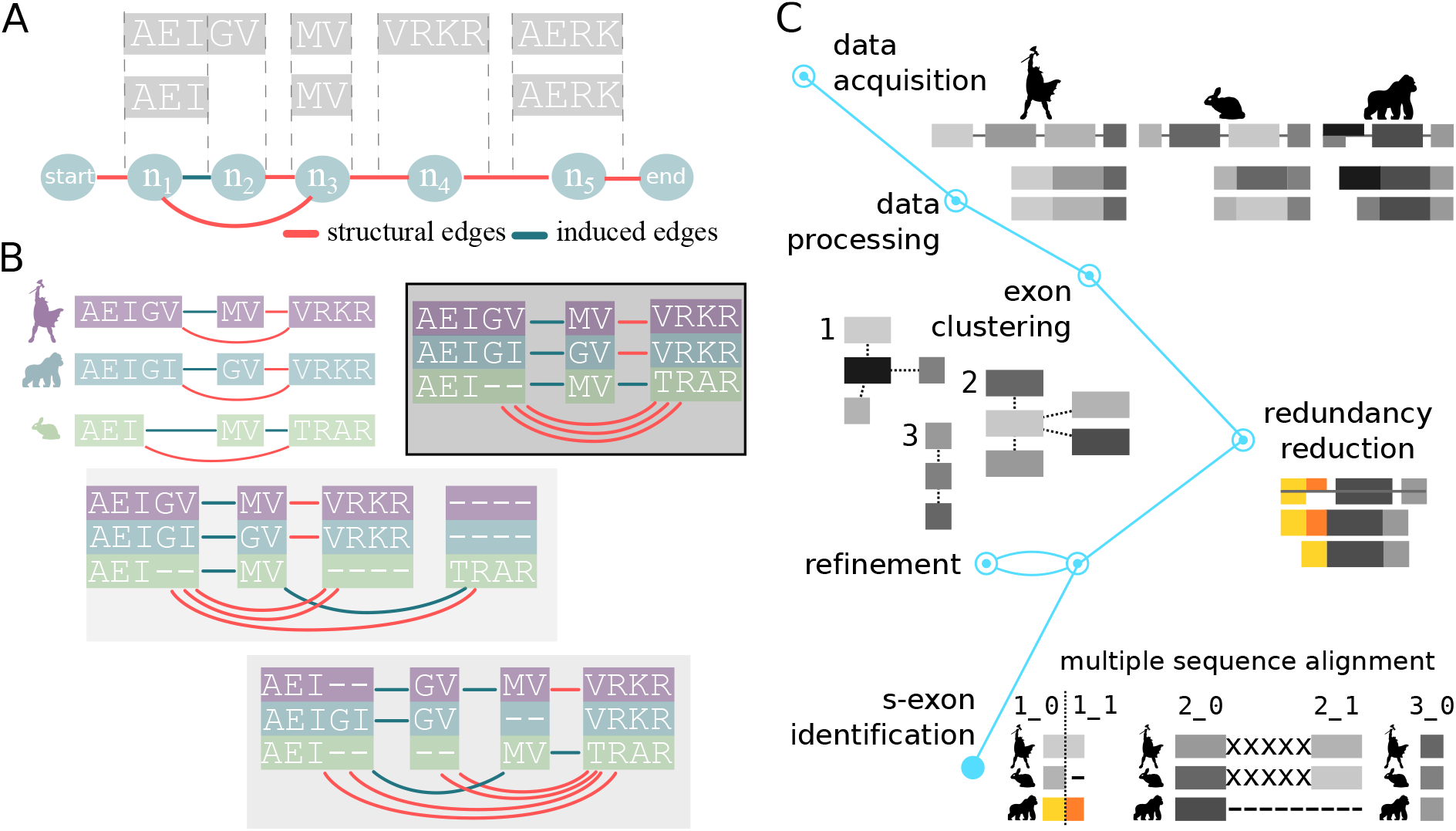
Principle of the method. **A**. Two transcripts are depicted, where each grey box represents a genomic interval and contains the corresponding protein sequence. Below, the minimal splicing graph (SG) is shown, with the nodes *n*_1_, *n*_2_…etc, corresponding to sub-exons. The *start* and *end* nodes are added for convenience. The structural edges in red correspond to some introns while the induced edges in green correspond to zero-base pair junctions. **B**. The top left panel shows a close-up view of three SGs corresponding to three orthologous genes coming from human, gorilla and rabbit. The information contained in these graphs can be summarised in one evolutionary splicing graph (ESG). Examples of ESG are depicted in the other panels, with the background grey tones indicating the ESG scores, the darker the better. The nodes in the ESGs represent *s-exons*, or multiple sequence alignments of exonic regions. Intuitively, the best-scored ESG shows at the same time compactness (parsimony) and good-quality alignments. **C**. Schematic workflow of ThorAxe pipeline. On top, the input genes and transcripts are displayed. The exons (grey boxes) are first clustered based on their similarities, then split into sub-exons to account for intra-species variability (boxes coloured in yellow and orange). Finally, the sequences belonging to each cluster are aligned and blocks in the alignments are identified to output a set of s-exons (*1_0*, *1_1*…). See **Supplementary Figure S1** for examples of intermediate and final outputs.

Our main contribution is to extend the definition of splicing graph to a set of orthologous genes *G* = {*G*_1_, *G*_2_,…, *G_n_*}. We define an evolutionary splicing graph (ESG) 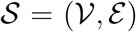 as a graph representing the whole transcript variability of *G*. Each 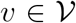 is a s-exon and represents a multiple sequence alignment (MSA) of sub-exons or sub-exons parts coming from different species. Each pair of nodes is linked by a set of edges deduced from the individual SGs. Among the many possible ways of building an ESG (**Fig. 1B**), one would like to find a representation as compact as possible and at the same time, conveying meaningful evolutionary information. To estimate both properties, we define the score of the ESG 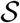 as

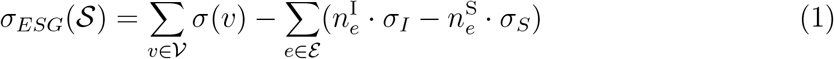

where *σ*(*v*) is the score of the MSA associated to the node *v*. *σ* can be for instance a consensus score or a sum-of-pairs score, and may additionally penalise very short MSAs (< 3 columns). 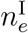 (resp 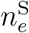) are the numbers of induced (resp. structural) edges in the multi-edge *e* with associated fixed penalty *σ_I_* (resp. *σ_S_*). In practice, we set *σ_I_* ≫ *σ_S_* to avoid small s-exons induced by ambiguous alignment columns in the MSAs. As an example, with a simple sum-of-pair scoring for *σ* and an induced edge penalty *σ_I_* = 4·*σ_S_*, the best-scored ESG in Figure 1B (framed and with the darkest background) comprises the smallest numbers of s-exons, induced edges and gaps. In general, determining the best-scored ESG is a NP-hard problem (see *Methods*).

Here, we provide a practical solution to construct a meaningful parsimonious ESG, given a set of input transcripts (**Fig. 1C**). Specifically, we have developed a heuristic procedure combining pairwise-alignment-based pre-clustering of exonic regions with cluster-specific MSAs complying with genomic constraints. The clustering step provides a coarse-grained partitioning of the sequence space that reduces the complexity of the MSA step. We set the parameters to obtain clusters small enough to produce high-quality MSAs but large enough to ensure a high species representativity of the s-exons. Then, we globally align all the sequences belonging to the same cluster using ProGraphMSA (Szalkowski, 2012). The latter allows better handling of AS-induced deletions and insertions than classical progressive alignment methods. Moreover, the genomic constraints help to disentangle orthology from paralogy relationships between similar sequences (**Supplementary Fig. S1**). Finally, we locally solve the problem exposed in Eq. 1 by re-aligning some sequences and by maximising the agreement between sub-exon boundaries across different species (**Supplementary Fig. S2**). The procedure is general enough to deal with very different genes (in terms of length, structure, degree of conservation, number of transcripts, etc.). We implemented it in the fully automated tool ThorAxe. All parameters are customisable by the user, enabling a rapid adaptation of the method to specific contexts and questions. Moreover, the output ESGs can easily be augmented and annotated with data coming from RNA-Seq experiments.

Our approach provides a double generalization. Firstly, we extend the definition of SG to the case of multiple species. Secondly, we provide a way to combine MSAs over structures with a partial order. In the following, we show that this formulation allows obtaining simple and meaningful representations for evolution in the context of AS.

### ThorAxe recapitulates known functional AS events

ThorAxe was tested on a curated set of 50 genes (16 families) across 12 species, from human to nematode (**Supplementary Table S1**). Within each gene family, several splice variants have been identified and associated to diverse protein functions (**Supplementary Table S2**). ThorAxe detected 93% of the functional AS events (ASEs) documented in the literature. All the detected ASEs are conserved in at least two species, and most of them are tissue-regulated (**Supplementary Table S3**).

As an illustrative example, we show the ESG computed for *CAMK2B* linker region on Figure 2A. Despite the very high complexity AS generates in this region, ThorAxe results are interpretable, meaningful and consistent with what has been reported in the literature. For instance, one can readily see that the shortest isoform lacking the linker has low evolutionary support (**Fig. 2B**, penultimate in the list). This is in line with recent findings emphasising the importance of the linker for regulating the protein activity (Bhattacharyya et al., 2020). Moreover, all the s-exons defined by ThorAxe are conserved in at least 7 out of 10 species (**Fig. 2B**). The smallest s-exon (*25_1*), of only one column, displays an alanine conserved from human to zebrafish. It corresponds to a well-documented internal splice site (Sloutsky and Stratton, 2020). Finally, the two documented functional ASEs (grey areas) are clearly identifiable on the ESG. This observation still holds true when scaling up to about 100 species (**Supplementary Fig. S3**). Furthermore, mapping RNA-Seq data onto the ESG (see *Materials and Methods*) revealed that the two events are tissue-regulated and that such regulation is evolutionary conserved (**Fig. 2C**). For instance, the alternatively spliced F-actin binding region comprised of the s-exons *15_0* and *15_1* is specifically expressed in the brain and muscles of primates and rodents (**Fig. 2C**, on the left).

**Figure 2:**
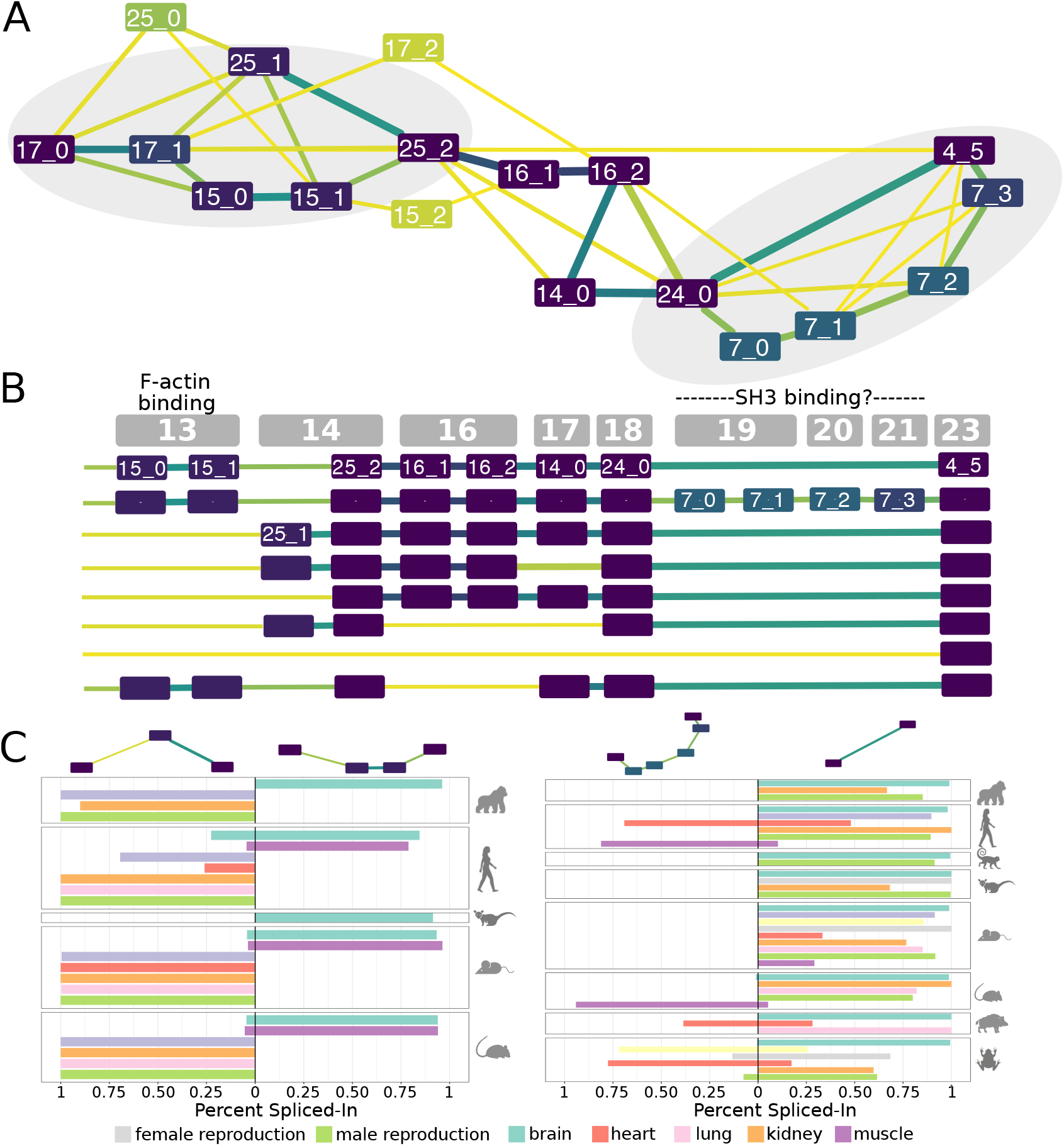
Transcript variability in the *CAMK2B* linker. **A.** Evolutionary splicing subgraph computed by ThorAxe starting from 63 transcripts annotated in 10 species. It corresponds to the region linking the kinase and hub domains of *CAMK2B*. The colours of the nodes and the edges indicate conservation levels, from yellow (low) to dark purple (high). Conservation is measured as the *species fraction* for the nodes (proportion of species where the s-exon is present) and as the *averaged transcript fraction* for the edges (averaged transcript inclusion rate of the s-exon junction). For ease of visualisation, the s-exons present in only one species were filtered out. The ASEs documented in the literature are located in the grey areas. **B.** On top, genomic structure of the human gene. Each grey box corresponds to a genomic exon (nomenclature taken from (Sloutsky and Stratton, 2020)). Below, list of human transcripts. All of them have been described in the literature, referred to as *β* (Bulleit et al., 1988), *β_M_* (Bayer et al., 1998), *β_e_* (Brocke et al., 1995), 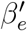 (Brocke et al., 1995), *β_e_*− (Cook et al., 2018), *α* (Bulleit et al., 1988), 7 (Wang et al., 2000) and 6 (Wang et al., 2000). The functional roles of some exons (Bayer et al., 1998; Khan et al., 2019) are given. **C.** Percent-Spliced In (PSI) computed from RNA-Seq splice junctions for the two documented ASEs. The two pairs of alternative subpaths depicted on top are also highlighted on panel A.

### ThorAxe summarises within and across-species variations at the human proteome scale

ThorAxe was further assessed on the whole human proteome (18,226 human protein coding genes, see *Materials and Methods*). The analysis performed across the same set of 12 species completed in less than 20 hours with 15 cores (**Supplementary Text S1**). One-to-one orthologs could be found in primates and mammals for almost three quarters of the human query genes, and also in amphibians and fishes for about half of them (**Supplementary Fig. S4**). On average, ThorAxe identified 26 s-exons per gene, but this number varies widely between genes (**Supplementary Table S4**).

S-exon conservation was measured by computing the *species fraction* (*F*), defined as the ratio between the number of species where a s-exon is present (number of sequences in the MSA) and the number of species considered (see *Materials and Methods*). The vast majority of the s-exons are either very lowly conserved or highly conserved (**Fig. 3A**). *Species-specific* s-exons, with only one sequence in the MSA, represent 40% of the ensemble (**Fig. 4A-B**, see nodes in yellow). They are often located at the N- and C-terminal extremities of the transcripts, and tend to display smaller and more variable lengths than the other *conserved* s-exons (**Supplementary Fig. S5**). About 80% of the latter are present in at least 80% of the species.

**Figure 3:**
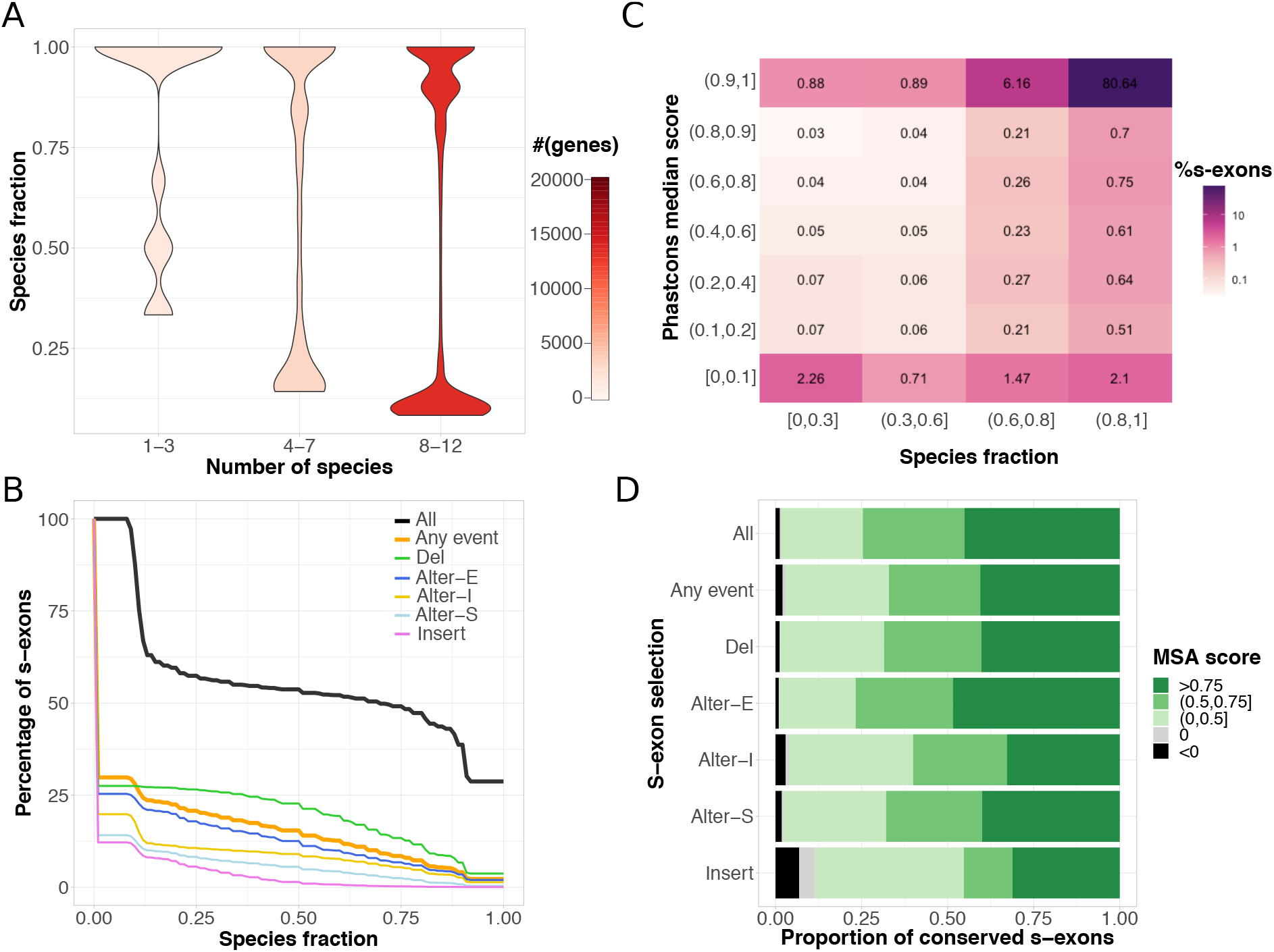
S-exon properties computed over the human genome. **A**. Distributions of the s-exon species fractions depending on the number of species. The red tones indicate the number of genes comprised in each distribution. **B**. Cumulative distributions of s-exon species fraction. On the y-axis we report the percentage of s-exons with a species fraction greater than the x-axis value. The different curves correspond to all s-exons (*All*), only those involved in at least an ASE (*Any event*), or only those involved in a specific type of event. *Alter-S*: alternative start. *Alter-I*: alternative (internal). *Alter-E*: alternative end. *Del*: deletion. *Insert*: insertion. **C**. Heatmap of the s-exon Phastcons median scores versus the s-exon species fractions. Only the s-exons longer than 10 residues and belonging to genes with one-to-one othologs in at least 8 species are shown. **D**. Proportions of conserved s-exons displaying very poor (negative score) to very good (score close to one) alignment quality. The MSA score of a s-exon is computed as a normalised sum of pairs. A score of 1 indicates 100% sequence identity without any gap. The proportions are given for different s-exon selections (same labels as in panel B).

**Figure 4:**
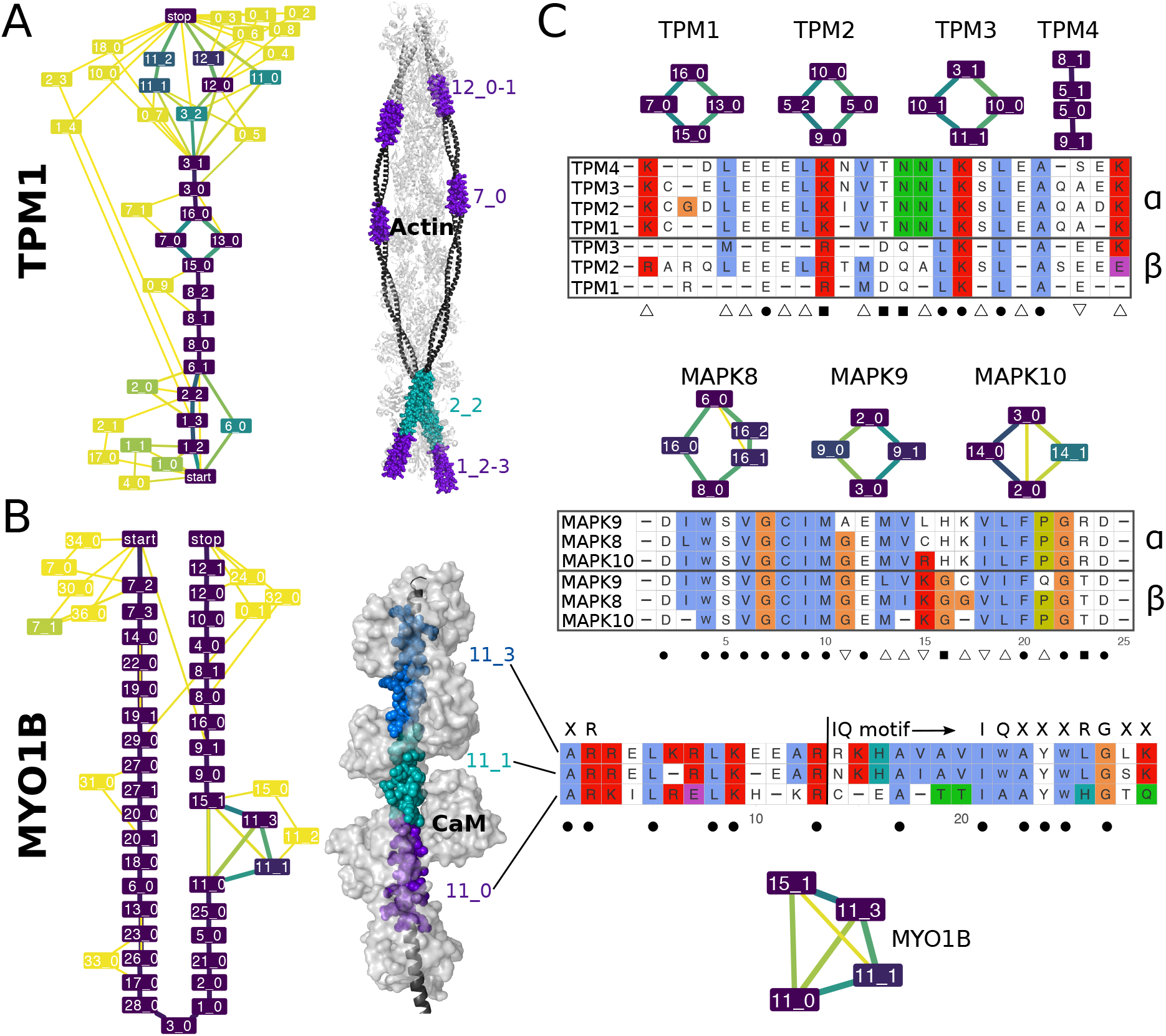
Examples of evolutionary conserved ASEs with in-gene paralogy. **A-B.** Evolutionary Splicing Graphs computed by ThorAxe (on the left) and the best 3D templates found by HHblits (on the right, PDB codes 2w49:abuv and 2dfs:H) for *TPM1* and *MYO1B*. On the ESGs, the colours indicate conservation levels, *species fraction* for the nodes and *averaged transcript fraction* for the edges (see *Materials and Methods*). The nodes in yellow are species-specific while those in dark purple are present in all species. The 3D structures show complexes between the query proteins (in black) and several copies of their partners (in light grey). The s-exons involved in conserved ASEs are highlighted with coloured spheres. **C.** S-exon consensus sequence alignments within a gene family (*TPM* on top, *MAPK* in the middle) or a gene (*MYO1B*, at the bottom). Each letter reported is the amino acid conserved in all sequences of the corresponding MSA (allowing one substitution). The colour scheme is that of Clustal X (Thompson et al., 1997). The subgraphs show the ASEs in which the s-exons are involved. The symbols *α* and *β* on the right indicate groups of s-exons defined across paralogous genes based on sequence similarity (see *Materials and Methods*). The symbols at the bottom denote highly conserved positions across the gene famil. Dot: fully conserved position. Square: position conserved only within each s-exon group. Upward triangle: position conserved in the *α* group only. Downward triangle: position conserved in the *β* group only. For *MYO1B*, the start and canonical sequence of the CaM-binding IQ motif are indicated. The motifs resulting from different combinations of the depicted s-exons are numbered 4, 4/5 and 4/6 in the literature (Greenberg and Ostap, 2013).

Almost a third of the s-exons are involved in some ASE (**Fig. 3B**). Those involved in deletions and insertions are the most and least conserved, respectively (**Fig. 3B**, green and pink curves). This is explained by the fact that ThorAxe detects ASEs as variations from a reference *canonical* transcript chosen for its high conservation and length (see *Materials and Methods*). The s-exons involved in internal alternative events and alternative starts tend to be more conserved than those involved in alternative ends (**Fig. 3B**, see the flatter gold and light blue curves compared to the blue one).

### The s-exons accurately measure sequence conservation

The s-exon conservation levels computed by ThorAxe are in very good agreement with estimates of evolutionary conservation deduced from whole genome alignments (summarized by Phastcons scores (Siepel and Haussler, 2005; Siepel et al., 2005), **Fig. 3C**). Moreover, for a significant proportion of s-exons, ThorAxe constructs MSAs with high median sequence similarity while genome-wide median conservation is very low (**Fig. 3C**, bottom right corner, and **Supplementary Fig. S6**). These s-exons are neither over-nor under-represented in ASEs.

Overall, almost half of the conserved s-exons are associated with very high quality MSAs, with a close-to-perfect percentage of identity (**Fig. 3D**). This proportion increases up to about 70% on the 50-gene set (**Supplementary Fig. S7**). The quality of the MSAs is lowered down for the s-exons involved in insertions, and to a lesser extent in internal alternative events (**Fig. 3D**). A very small proportion (about 1%) of s-exons, typically comprised of very few positions (**Supplementary Fig. S8**), display very poor (negative) MSA scores.

### The comparison of similar s-exons unveils functionally relevant signatures

ThorAxe allows exploring how function diversification may arise through the alternative usage of similar sequences within and across genes. We illustrate the power of the approach on three gene families (**Fig. 4**, and see also **Supplementary Table S2-3**). We focus on a set of ASEs involving two or more s-exons highly conserved in evolution and whose consensus sequences share some similarity. The origin of the ASEs can be traced back to the ancestor common to mammals, amphibians and fishes.

The first example is given by the tropomyosin family (**Fig. 4A,C**), whose members (*TPM1,2,3,4*) serve as integral components of the actin filaments forming the cell cytoskeleton. Several conserved ASEs are observed on the ESGs, and the alternatively spliced s-exons directly interact with actin (**Fig. 4A**). Among them, the internal mutually exclusive pair displays high sequence similarity. The s-exon consensus sequences reveal a strong conservation signal across species and between paralogous genes (**Fig. 4C**, on top, see *α* and *β* groups). Fourteen specificity-determining sites (SDS) can be identified (**Fig. 4C**), 11 of type I (triangles) and 3 of type II (squares). SDS are key positions with specific conservation patterns, and they play a role in diversifying protein function in evolution (Chakraborty and Chakrabarti, 2015). Given two groups, here *α* and *β*, type I SDS are conserved in one group and variable in the other one, indicating different functional constraints between the groups. For instance, position 24 is occupied by a glutamate in all the s-exons from the *β* group, while it is variable in the *α* group. Type II SDS are conserved in both groups but each group displays a different amino acid. This is the case of positions 14 and 15, where the Thr-Asn couple of the *α* group is replaced by Asp-Gln in the *β* group. The identified SDS may be responsible for the differences observed in actin filaments formation, mobility and myosin recruitment ability between the isoforms (Pathan-Chhatbar et al., 2018).

The c-Jun N-terminal kinase family (*MAPK8,9,10*) gives another example with even higher sequence identities (**Fig. 4C**, in the middle). Eight SDSs of type I and 2 of type II are identifiable. In particular, three positively charged residues, His, Lys and Arg, in positions 16, 17 and 23 are specifically conserved in the *α* group, while the *β* group is characterised by Lys, Gly and Thr in positions 15, 16 and 23. These observations are in line with our previous study highlighting differences in the dynamical behaviour of these residues (Ait-hamlat et al., 2020). Such differences may play a key role in determining the isoform-specific substrate selectivities (Waetzig and Herdegen, 2005).

As a third example, Myosin-1B is characterised by evolutionary conserved insertions of two very similar s-exons that also resemble the constitutively expressed s-exons located upstream in the ESG (**Fig. 4B**). Each s-exon overlaps with two calmodulin(CaM)-binding so-called IQ motifs, such that different s-exon combinations lead to different binding motifs. The s-exon consensus sequences share almost 50% identity (**Fig. 4C**, at the bottom). Compared to the motif’s canonical form (IQXXXRGXXXR) (Houdusse et al., 1996; Bähler and Rhoads, 2002), they all lack the glutamine in the IQ residue pair and the arginine in the RG pair. These differences could explain their lower affinity compared to the constitutive s-exons (Greenberg and Ostap, 2013).

### Alternative usage of similar sequences is not a rare phenomenon

At the human genome scale, we identified 2 190 genes with evidence of evolutionary conserved alternative usage of similar exonic sequences (**Fig. 5**). This represents about 12% of the protein-coding genome. Each of these genes comprises two or more s-exons sharing significant sequence similarity, and possibly corresponding to several case scenarios, as described below.

**Figure 5:**
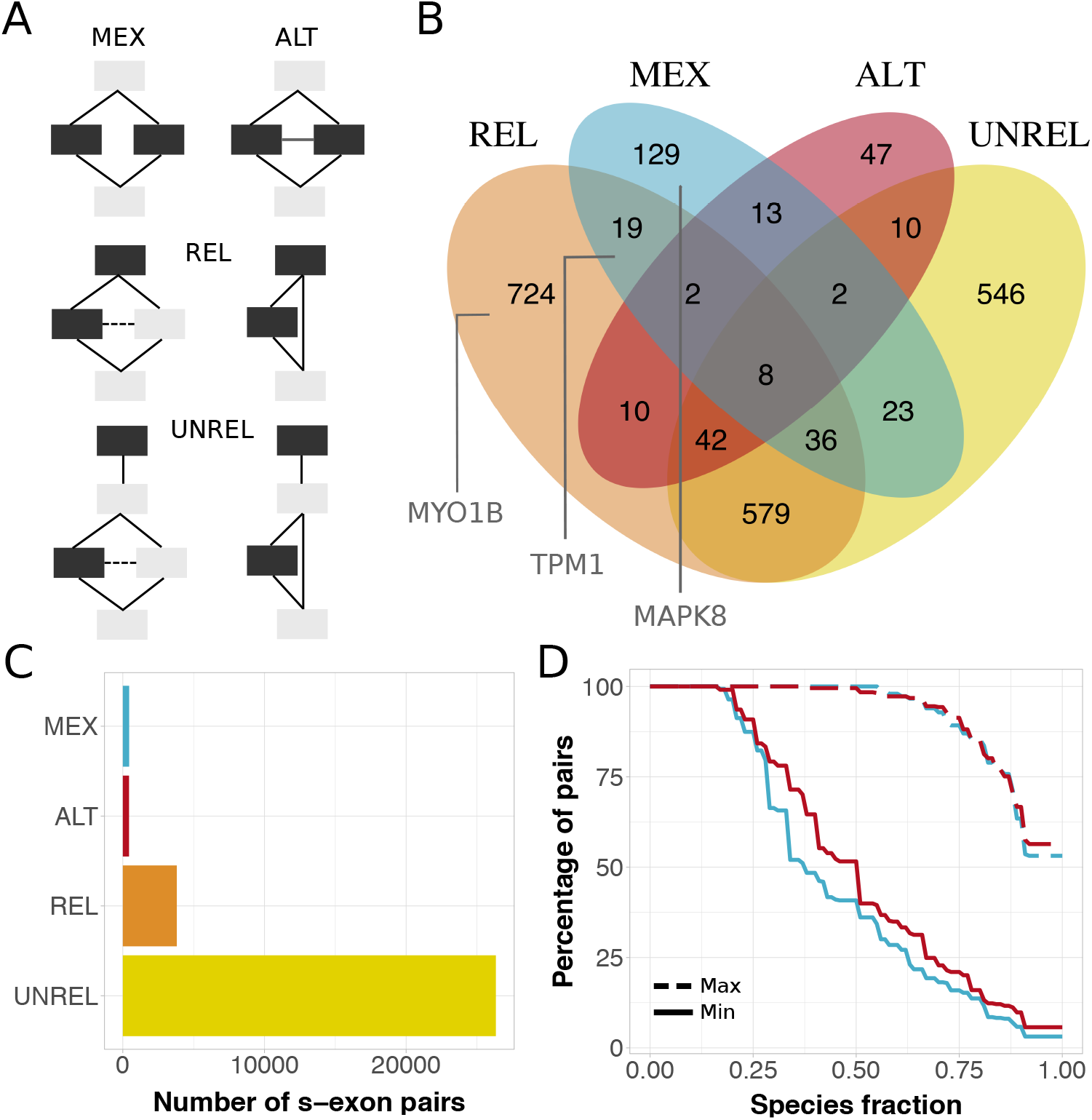
Alternative usage of similar s-exons. **A.** Evolutionary splicing subgraphs depicting alternative usage different scenarios. The detected s-exon pairs are colored in black. MEX: mutual exclusivity. ALT: alternative (non mutually exclusive) usage. REL: one s-exon is in the canonical or alternative subpath of an event (of any type), while the other one serves as a “canonical anchor” for the event. UNREL: one s-exon is in the canonical or alternative subpath of an event (of any type), while the other one is located outside the event in the canonical transcript. Each detected pair is assigned to only one category with the following priority rule: MEX>ALT>REL>UNREL. **B.** Venn diagram of the genes containing similar pairs of s-exons. The genes shown in Figure 4 are highlighted in the corresponding subsets. **C.** Proportions of s-exon pairs detected in the four categories. **D.** Cumulative distributions of s-exon conservation. On the y-axis we report the percentage of s-exon pairs with species fraction greater than the x-axis value. The solid (resp. dashed) curve corresponds to the highest (resp. lowest) species fraction among the two s-exons in the pair. We report values only for the MEX (in blue) and ALT (in red) categories.

Out of the 31 031 similar s-exon pairs we detected, 446 pairs are mutually exclusive (**Fig. 5A**, MEX). This case scenario highlights the exclusive usage of one or the other version of a protein region. Another 438 pairs are alternatively used without mutual exclusivity (**Fig. 5A**, ALT). In about half of the s-exon pairs classified as MEX or ALT, one of the s-exons is conserved in all studied species, while the other one is conserved in more than half of them (**Fig. 5D**). This is in line with a previous study reporting a high evolutionary conservation for mutually exclusive homologous exon pairs (Abascal et al., 2015). In 3 813 pairs, one s-exon is included in the canonical or alternative subpath of an ASE, while the other one serves as a “canonical anchor” for the ASE (**Fig. 5A**, REL). This highlights the AS-induced modulation of the number of non-identical consecutive copies of a protein region. The remaining 26 334 pairs correspond to cases where the s-exons are not found in the same ASE (**Fig. 5A**, UNREL). Hence, the modulation occurs between non-consecutive regions. The full lists of detected s-exons, with detailed information about their lengths, categories, and involvement in ASEs are given in Supplementary Tables 5-7.

The resource we provide here overlaps well with a previously reported manually curated set of 97 human genes with mutually exclusive homologous exon pairs (Abascal et al., 2015). It extends it by one order of magnitude and represents a more diverse range of AS-mediated relationships between similar protein regions. The proteins we identified tend to be involved in cell organisation and muscle contraction (cytoskeleton, collagen, fibers…etc), and in inter-cellular communication (**Supplementary Fig. 9**). The MEX-containing protein subset is specifically enriched in transporters and channels. Overall, nebulin gives the most extreme example, with 8 802 detected s-exon pairs. This giant skeletal muscle protein has evolved through several duplications of nebulin domains, and a definition of pertinent nebulin evolutionary units was proposed (Björklund et al., 2010). These units correspond to parts of exons, in line with ThorAxe s-exons MSAs.

### Comparison with other methods

ThorAxe produces longer and higher-quality s-exon MSAs compared to other strategies that would omit the clustering step or rely solely on global multiple sequence alignment (**Fig. 6A-B**). Specifically, the clustering step helps to improve the s-exon sequence identities (**Fig. 6B**, compare blue and orange boxes) by reducing the space of sequences to align. The final local optimization step increases the lengths of the s-exons (**Fig. 6A**, compare blue and red boxes) by minimising sub-exon boundaries violations. Compared to the Reciprocal Best Blast Hit (RBBH) method, ThorAxe achieves a higher sub-exon coverage (**Fig. 6C**). This result highlights the importance of the one-to-many sub-exon matching between species. Exons may undergo truncation or elongation in the course of evolution, and thus we do not expect a one-to-one relationship between them across a pair of species. ThorAxe increased coverage is not at the expense of sequence identity (**Supplementary Fig. S10**). Compared to Ensembl Compara (Herrero et al., 2016), ThorAxe extends the applicability of exon orthology detection to all the species for which annotated transcripts are available. Moreover, it produces higher quality alignments by considering the translated sequence.

**Figure 6:**
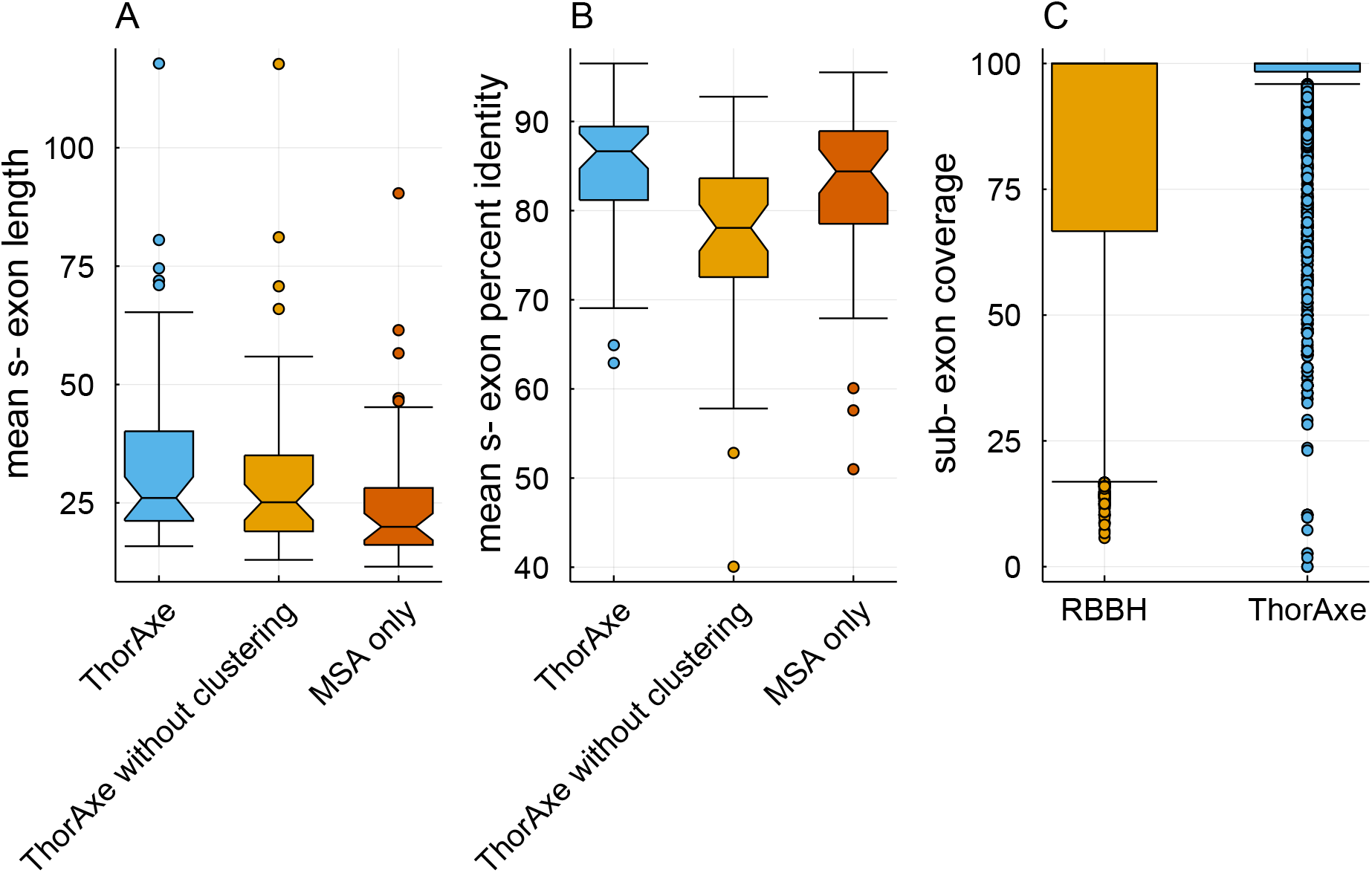
Comparison of ThorAxe performance against other strategies. **A-B.** Comparison between ThorAxe (in blue), a strategy combining MSA and ThorAxe refinement procedure (in orange), and a strategy relying solely on MSA (in red). In all cases, the MSAs are produced using ProGraphMSA (Szalkowski, 2012). **A.** Distributions of s-exon lengths averaged over each gene. **B.** Distribution of s-exon MSA percent identities averaged over each gene. **C.** Distributions of sub-exon coverages obtained from RBBH (in orange) and ThorAxe (in blue). The ensemble of genes considered are those from the curated set.

## Discussion

We have presented a novel method to describe transcript variability in evolution. It produces compact representations allowing for a direct assessment of transcripts and AS events evolutionary conservation. It can deal with diverse genes, long evolutionary times, and potentially very many species. A crucial aspect of this work distinguishing it from previous developments is the generalisation of the notions of exon and splicing graph (Heber et al., 2002) by adding an evolutionary dimension. We provide a practical solution for defining the minimal building blocks (the s-exons) for reconstructing a set of transcripts across several species, which is the first and necessary step for inferring evolutionary scenarios explaining AS-induced protein function diversification (Ait-hamlat et al., 2020). Moreover, we provide new and more accurate estimates of the conservation of AS for protein-coding sequences across many species. We have also shown that the framework presented here is useful to decipher the complexity of AS-induced sequence variations toward the identification of specificity-determining sites (Chakraborty and Chakrabarti, 2015). This finding calls for the systematic detection of sequence conservation patterns through the prism of the s-exon.

Our data structure is reminiscent of current developments on pangenome graphs. However, pangenome approaches keep track of variations across a population, whereas we highlight conservation across species in the context of AS. To the best of our knowledge, we are the first to do it. Effectively, we consider a *pan-transcriptome* across multiple species. As a consequence, we do not need to rely on a central species and project the transcripts on it. In the analysis conducted here, human was taken as a reference only to find orthologous genes in other species.

The constructed evolutionary splicing graphs depend on gene annotations that may be partial, incomplete, or erroneous (Salzberg, 2019). To ensure the high quality of the input data, we proceed through a careful pre-processing step accounting for transcript support. Moreover, to cope with possible errors, our algorithm is designed to handle peculiar variations occurring in only one transcript or one species. We have shown that these variations are associated with low conservation levels and that it is easy to segregate and filter them out for further analysis. Finally, one can easily complement the ESG by information coming from raw RNA-Seq data, allowing the discovery of new events and the assessment of transcript tissue regulation.

We have also collected two medium-size sets of genes that can be useful to the community. The first one is relevant to the study of AS functional impact. It will serve as a reference set for future studies. The results we compiled (ESGs deduced from transcript annotations (Yates et al., 2016), enriched with RNA-Seq data (Komljenovic et al., 2018)) can help to gain insight into essential biological processes such as muscle contraction and transport. The second one opens avenues to in-depth studies of the alternative usage of repeats in proteins. We have shown that this mechanism is not a rare phenomenon in the human proteome and that it is of ancient evolutionary origin.

Although we found that our heuristic produces interpretable results consistent with the literature and is robust enough for large-scale application, future work would likely benefit from the development of more accurate approximations of the general problem stated here. Another direction for the future is to expand the application field to transcriptomes coming from patients or human populations. In the coming years, we expect a tremendous increase in the available transcriptomic data, including transcriptome annotations generated by long-read sequencing technologies (Byrne et al., 2019). Methods addressing the complexity of these data will become instrumental in understanding the evolution of a disease, *e.g.*, cancer, and the phenotypic variability among human populations and individuals (Park et al., 2018; Lonsdale et al., 2013).

## Methods

### Datasets

#### AS-dedicated gene set

We collected a set of 50 genes, representing 16 families, where AS produces functionally distinct protein isoforms (**Supplementary Table S1**). Within each family, the biochemical activities of several isoforms have been characterised. The set contains single-domain genes as well as multi-domain ones, ranging from less than 200 to 2000 residues. It includes some kinases, some receptors, some RNA-binding proteins, a transcription factor, and a significant portion of proteins involved in the formation of the cell cytoskeleton, in muscle contraction and in membrane trafficking.

#### Human proteome

We considered the ensemble of 19,976 protein coding genes comprised in the human genome and their one-to-one orthologs from gorilla, macaque, opossum, rat, mouse, cow, boar, platypus, xenopus, zebrafish and nematode. We downloaded the corresponding gene annotations from Ensembl (Yates et al., 2016) release 98 (September 2019). The download was successful for 18,241 genes. Among those, 14 genes did not have any good quality transcripts (see below for the criteria) and 1 gene displayed an error in the gene tree, leading to a total of 18,226 valid genes. We should stress that a small fraction of these genes (3%) do not have any valid human transcript.

### Definitions

#### Splicing Graph (SG)

Given a gene *G_i_*, and its annotated transcripts described as sorted lists of genomic intervals, we define a *splicing graph* (*SG*) for *G* as the directed graph 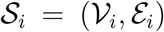 (**Fig. 1A**). Each node 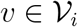 is identified by (*v_s_*, *v_e_*, *v_f_*), where [*v_s_*, *v_e_*] is the genomic interval covered by *v* and *v_f_* is its reading frame. There is a node *v* in 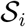 if the corresponding coordinates and frame occur in at least one transcript. There is an edge in 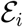 from *v* to *v*′ if the corresponding coordinates and frames are consecutively transcribed. Edges are classified either as *structural* if *v* and *v*′ are separated by an intron or *induced* by the nodes’ genomic boundaries otherwise. Finally, a *start* and *end* nodes are added and act respectively as the least and the greatest element for the partial order induced on the graph. Note that, due to the partial order on genomic intervals and the colinearity of transcripts in the genome, the graph 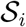 is directed acyclic.

In principle, there are many possible splicing graphs, depending on how the nodes are defined. The *minimal splicing graph* is the unique SG with the smallest number of nodes, such that each one of the input transcripts is represented by a path. When not stated explicitly we will always refer to the minimal splicing graph.

#### Sub-exon

We define sub-exons as nodes in a minimal SG. They are the minimal building blocks for the transcripts observed in a given species. As an illustrative example, let us consider the gene SNAP25 from gorilla (**Supplementary Fig. S1B**). The two first transcripts, TRX1 and TRX2, start with the same exon (`MAE…ADE`, in yellow) comprised of 24 residues. The third transcript, TRX3, starts with another longer exon (`MAE…GRE`, in yellow and orange) comprised of 33 residues and overlapping with the previous one. In that case, we define two sub-exons, one of 24 residues (`MAE…ADE`, in yellow) and another of 9 residues (`VRS…GRE`, in orange).

#### Evolutionary Splicing Graph (ESG)

We extend the definition of the SG to a set of orthologous genes *G* = {*G*_1_, *G*_2_,…, *G_n_*} (**Fig. 1B**). Our aim is to construct a graph summarising transcript information from 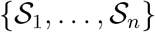 together with evolutionary conservation by means of an *evolutionary splicing graph* (*ESG*) 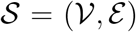. Although some cycles may appear in 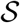 due to exon switching, the graph will be directed acyclic in most cases. Each 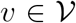 is formally described by a list 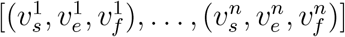 of genomic coordinates and reading frames for each of the species (some elements can be empty). In practice, we will be interested in the corresponding multiple sequence alignments (MSAs) of the translated sequences. 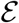 is composed of multi-edges, *i.e.*, sets of edges deduced from the individual SGs. Each multi-edge can be either purely structural, purely induced or hybrid, whenever several genomic exons are joined in some of the species.

As for the individual SGs, there are multiple ways of constructing 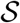. Our goal is to minimise the number of nodes while maximising the overall sequence similarity of the MSAs associated to each node. We formalise the problem using a scoring function (Eq. 1). A *minimal evolutionary splicing graph* is simply a graph with maximal score. Note that there may exist several minimal ESGs for a given gene.

#### S-exon

We define s-exons as nodes in a minimal ESG. They are the minimal building blocks for the transcripts observed in a set of species. Let us consider again the example of SNAP25, studied across eight species (**Supplementary Fig. S1C-D**). Each s-exon is represented by a MSA, where each sequence comes from one species and corresponds to a sub-exon or a part of a sub-exon. The exonic sequences belonging to the same s-exon are supposed to be orthologous. A species-specific s-exon comprises only one sequence (coming from one species), while a conserved s-exon comprises at least two sequences (coming from two species).

### Complexity of the problem: determining a minimal ESG is NP-Hard

To illustrate the complexity of determining a minimal ESG, let us consider a case example with *n* input transcripts observed in *n* species (*i.e.*, 1 transcript per species). Moreover, since the problem is theoretically independent of the penalties *σ_I_* and *σ_S_*, an algorithm that would solve it in the general case would also be valid for *σ_I_* = *σ_S_* = + ∞. In this scenario, a minimal ESG has no edge and maximises the sum-of-pair-score *σ*. Thus, the problem of building a minimal ESG is equivalent to solving the problem of multiple sequence alignment with sum-of-pair-score *σ* on the *n* input transcripts. Since the *n* input transcripts can be any string (over the amino acid alphabet), and finding a multiple sequence alignment of any string with sum-of-pair-score is NP-hard (Wang and Jiang, 1994), it follows that finding a minimal ESG is NP-hard.

### Detailed description of ThorAxe algorithm and parameters

#### a- Data acquisition and pre-processing

ThorAxe takes as input a gene name and a list of species (**Fig. 1C**). First, the gene tree, the transcripts annotated as protein coding and the corresponding exons (genomic coordinates, sequences and phases) are retrieved from Ensembl, for the query gene and its one-to-one orthologs in the selected species. Second, a quality check procedure is applied to remove incomplete transcripts and transcripts with low level of support (TSL value lower than 3, adjustable by the user). Third, the DNA sequences of the retained transcripts are translated into amino acid sequences. To avoid redundancy, transcripts or exonic regions leading to the same protein sequence are merged.

#### b- Pairwise-alignment-based exon pre-clustering

Our initial building blocks are the exons annotated in Ensembl, for each gene within each species. The first step of our algorithm is to partition the exon space by grouping the exons sharing some similarity. The similarity between two exons is estimated by determining their optimal local alignment using the Smith-Waterman algorithm (Smith and Waterman, 1981). To reduce the computational burden of performing an all-to-all comparison, we implemented a modified version of the Hobohm I algorithm (Hobohm et al., 1992). The exons are first ordered from the longest to the shortest one. Then, at each step of the algorithm, we compare the current exon *i* with the following (smaller) exons *j*, with *j* > *i*. If exon *j* shares more than *id_cut_*% sequence identity with exon *i* and is covered at more than *cov_cut_*% by the alignment, then the two exons are assigned to the same cluster. By default, *id_cut_* = 30 and *cov_cut_* = 80, and these parameters can be adjusted by the user. Setting the identity threshold to a relatively low value ensures that homology is detected across many species. The clustered exons are progressively removed from the initial set, such that they will not be considered in the next iteration of the algorithm for the comparisons. We modified the algorithm to be more stringent on this latter criterion. Specifically, if exon *j* gets assigned to the same cluster as exon *i* but shares less than (*id_cut_* + 30)% sequence identity with *i*, then it remains in the set. It will be compared against the other exons in the next iteration, and will migrate from one cluster to another if we find a better match. Please note that some clusters may contain only one exon.

#### c- Redundancy reduction

Within each species, ThorAxe systematically identifies sets of overlapping exons, and replaces them by non-redundant distinct sub-exons. This step relies only on the genomic coordinates of the exons and does not require aligning the exonic sequences. It is performed after exon clustering, since dealing with sub-exons at this early stage would add some unnecessary complexity by augmenting the number of comparisons and the ambiguity associated with small sequences.

#### d- MSA-based s-exon identification

To determine a mapping of exonic sequences between different species, we rely on global multiple sequence alignment. We use the previously identified exon clusters to simplify the task. Specifically, for each cluster, we generate a MSA comprising *n* sequences, where *n* is the number of species with at least one exon in the cluster. Each sequence is a chimeric construct built by concatenating the sub-exons in the cluster coming from a given species, according to their genomic coordinates. A padding sequence of “X” is introduced between two sub-exons if they are never observed together in any transcript. This padding is used to enforce separating them in the MSA. To align the sequences, we use the graph-based progressive alignment method ProGraphMSA (Szalkowski, 2012). We chose this method because its graph-based representation of protein sequences allows recording the whole history of indel events along the guide tree. This framework proved better suited to deal with AS-induced insertions and deletions than classical progressive alignment methods (Szalkowski, 2012). Finally, the s-exons are identified as continuous blocks in the MSA, delimited by the sub-exon boundaries. In brief, we scan the MSA from left to right and create a new s-exon whenever there is a change of sub-exon in at least one sequence. This ensures that the identified s-exons can be used as building blocks to reconstruct any transcript in any species from the input data. A detailed description of the s-exon identification algorithm is given in Supplementary Text S1 (see Algorithm 1).

#### e- S-exon refinement

To improve the definition of the s-exons, we apply a refinement procedure locally optimising the chimeric MSAs. Firstly, we migrate lowly-scored sub-exons from one MSA to another. By default, we consider that a sub-exon is poorly aligned if it shares less than 30% sequence identity with all the other sequences against which it is aligned. ThorAxe algorithm aligns the problematic sub-exon to sequences from the other clusters, and rescue it in case a better match is found. Otherwise, a new species-specific s-exon is created containing only the sub-exon sequence. Secondly, we minimise the number of very small s-exons, comprising only 1 or 2 columns. Typically, these s-exons arise from inconsistencies between the sub-exon boundaries in the different species of the MSA (**Supplementary Fig. S2**). To minimise such inconsistencies, ThorAxe algorithm shifts the very small sequence stretches isolated by gaps to re-group sub-exons (**Supplementary Fig. S2**, bottom left panel). In addition, it disintegrates the 1-column s-exons by re-assigning their residues to the neighbouring s-exons, whenever this is consistent with sub-exon boundaries (**Supplementary Fig. S2**, right panels).

#### f- ESG construction and annotation

Once the s-exons have been identified, building the ESG is straightforward. To ease visualisation, ThorAxe builds a compressed ESG, in which the multi-edges are represented by single edges. Moreover, it annotates the nodes and the edges of the graph with evolutionary information and summary statistics. For a node *v*, let us denote as *n*(*v*) the number of species where *v* is present, *nt*(*v*) the number of transcripts containing *v*, *nt_s_* the number of transcript of species *s*, *nt_s_*(*v*) the number of transcripts from species *s* containing *v*, and *nt* the total number of input transcripts (*nt* = ∑_*s*_ *nt_s_*). We consider three conservation measures:

- the *species fraction*, 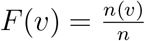
- the *transcript fraction*, 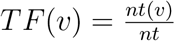
- *averaged transcript fraction*, 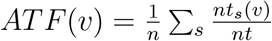.

These measures can be expressed in the same way for a given edge *e*, or for a given path in the graph. The *species fraction* indicates in how many species a node/edge/path is observed, the *transcript fraction* indicates in how many of the input transcripts the node/edge/path is included and the *averaged transcript fraction* reflects the average transcript usage of the node/edge/path. The ESGs were visualised with Cytoscape V.3.7.2 (Shannon et al., 2003).

#### g- AS event detection

The AS events are defined as variations detected between each input transcript and a *canonical* transcript serving as reference. Ideally, the canonical transcript should be well represented across species. Hence, to decide which of the input transcripts should serve as reference, we look at the species fraction and the averaged transcript fraction of its edges. More specifically, we pick up the transcript *t* (described by the list of its edges) maximising three variables, in the following priority order: (1) min_*e*∈*t*_ *F*(*e*), (2) min_*e*∈*t*_*ATF*(*e*) and (3) 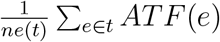, with *ne*(*t*) the number of edges in *t*. We should stress that this combination yielded the best results among all possible combinations of nine measures (minima, averages and sums of *F*, *TF* and *ATF*) on our curated set. If more than one candidate transcripts are found, we retain the longest (in residues) and highest-quality (based on TSL) one. To detect AS events, we enumerate all the pairs of maximal subpaths that do not share any s-exon, where one subpath comes from the canonical transcript and the other one from some input transcript (**Supplementary Text S1**, Algorithm 2). By default, the ASEs are detected on a reduced version of the ESG, where the edges supported by only one transcript have been removed.

#### h- Outputs and interfacing with other tools

In addition to the annotated ESG, ThorAxe outputs the list of input transcripts described as collections of s-exons (where each s-exon is designated by a symbol) and the gene tree representative of the selected species. These data can directly serve as input for PhyloSofS (Ait-hamlat et al., 2020), toward the reconstruction of transcripts’ phylogenetic forests. ThorAxe may also be easily interfaced with other tools requiring the same type of input.

### Calculation of MSA scores

The quality of the s-exon MSAs was assessed using a sum-of-pair score, with *σ_match_* = 1, *σ_mismatch_* = −0.5 and *σ_gap_* = 0. The computed raw scores were normalised by dividing them by the maximum expected values. Hence, the final score of the s-exon represented by the node 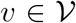 is expressed as,

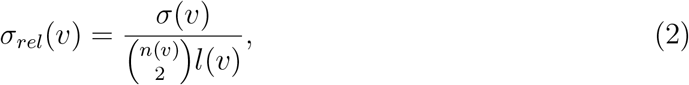

where *σ*(*v*) is the raw MSA score, *n*(*v*) is the number of species where the s-exon is present and *l*(*v*) is the length of the s-exon, computed as,

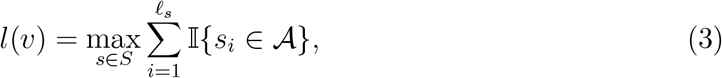

where *S* is the set of sequences comprised in the MSA, *l_s_* is the length of the aligned sequence *s*, and 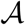 is the 20-letter amino acid alphabet (*e.g.*, excluding gap characters).

### Detection of similar pairs of s-exons

To systematically identify protein sequences sharing some similarity, we performed an all-to-all comparison of the s-exons identified by ThorAxe for each human protein coding gene. Specifically, we created a profile hidden Markov model (HMM) from each s-exon MSA using *hhmake* from the HH-suite (Steinegger et al., 2019). No filtering was applied on the s-exon sequences, and the maximum proportion of gaps in a MSA column to be considered for match states was set to 50%. For each gene, we globally aligned each HMM against all the others using hhalign. We considered two s-exons to be similar if the p-value associated to their HMM alignment was lower than 0.001. In total, we detected 150 020 s-exon pairs found in 10 814 genes. We further restricted this set to s-exons present in more than one species and involved in at least an ASE. In addition, we removed the s-exon pairs detected based on HMM alignments smaller than 5 positions, and the pairs where the two s-exons do not have any species in common. Finally, we excluded the pairs where none of the s-exons is included in the canonical transcript, and those where the two s-exons always co-occur in the same subpath (canonical or alternative). Indeed, these pairs do not inform us about the differential usage of similar sequences. These filters reduced the number of s-exon pairs to 31 031 and the number of genes to 2 190. Among those, we retrieved 90% of a previously reported set of 97 genes comprising pairs of mutually exclusive homologous exons (Abascal et al., 2015). Four of the missing genes (ACSL6, PPAP2A, U2AF1, UGT1A8) were not in the set of 18,226 human protein coding genes treated by ThorAxe. One of them (CYP4F3) did not display any ASE. In five other genes (H2AFY, HNRNPK, ITGA3, RBM4, SLC39A14), no significant similarity was detected between the mutually exclusive sequences reported in (Abascal et al., 2015).

### RNA-Seq analysis

We used the Bgee database (Komljenovic et al., 2018) to select a relevant set of 37 RNA-seq experiments over our 12 species of interest. The corresponding raw sequences were then downloaded from the Sequence Read Archive (SRA-NCBI) (Leinonen et al., 2010) (total: 877 libraries), and aligned to their respective genome versions with the STAR aligner (Dobin et al., 2012). The resulting BAM files were used to update the reference gene annotation of some species (see below). Whippet (Sterne-Weiler et al., 2018) was then run on the raw sequences for ASE quantification.

#### Data preparation

We assessed read libraries strandedness and quality with the RSeQC package (Wang et al., 2012). To obtain the strand information from the BAM files we used the function *infer experiment*. We determined the mapping quality with the function *bam stats* and kept only the files containing more than 75% of high quality mapped reads. The selected libraries were then merged for all species and experiments using SAMtools (Li et al., 2009).

#### Whippet annotations

Whippet analysis unfolds in two steps. First, it builds a species-specific index using GTF annotations and, optionally, BAM files. We used both types of files for macaque, rat, cow, platypus, xenopus, and zebrafish. For human, gorilla, boar, opossum and nematode, we used only GTF annotations. Indeed, the BAM files of gorilla, boar and nematode were of low quality and/or non-stranded. Hence, following Whippet developers’ recommandations, we did not use them for indexing. Moreover, another recommandation is that BAM files associated with well-annotated genomes should not be considered. This excluded the BAM files of human and mouse. Finally, the chromosomes of opossum were too long to be handled by SAMtools, and hence its BAM files were also ignored. Secondly, Whippet detects and quantifies the ASEs supported by the reads. Note that it does the quantification without aligning the reads. We used the function *whippet-quant.jl* for computing Percent-Spliced In (PSI) values (Venables et al., 2008; Katz et al., 2010). The function is suitable for treating both single-end and paired-end reads.

#### Extraction of splice junctions

The splice junctions obtained from Whippet were mapped on the ESGs computed by ThorAxe using genomic coordinates and strand information. For each splice junction, we computed the normalised sum of mapped reads to cope with sequencing depth variations between the different experiments.

#### Mapping between ThorAxe and Whippet nodes

We created a mapping between the s-exons identified by ThorAxe and the exonic regions represented by the nodes in Whippet. Let us remind that ThorAxe s-exons are defined across species while Whippet nodes are species-specific. Within each species and for each s-exon, we looked for matching Whippet nodes. We identified three main case scenarios (**Supplementary Fig. S11**): (*i*) one perfect match found, (*ii*) partial match(es) found at the 3’- and/or 5’-end, or (*iii*) no match found. Case (*i*) corresponds to a one-to-one non-ambiguous mapping. For case (*ii*), if two partial matches were found, one at each end, we defined a one-to-many mapping for the s-exon. If only one partial match was found, we inferred an overlap between a Whippet node and one of the s-exon extremity. Then, if the matching Whippet node was identified as a partial match for another s-exon at the other extremity, we inferred a many-to-one mapping. Otherwise, it meant a Whippet node corresponded to some s-exon(s) and some contiguous unannotated region. For dealing with case (*iii*), we re-ran the search with relaxed matching constraints. Specifically, if the s-exon was longer than 100 bp, its start and end coordinates were allowed to differ by 25% of its length, compared to the Whippet matching node(s). If the s-exon was shorter, then the allowed variation was set to 33% of the s-exon length. Finally, the PSI value of a s-exon in a given tissue from a given species was computed as the mean PSI value of its matching Whippet nodes.

#### Splice graph annotation and ASEs quantification

An edge in the ESG was considered as supported by RNA-Seq data in a given species if the normalised mean of mapped reads computed for the corresponding junction was higher than 1*e*− 07. For the nodes, we set up a PSI threshold of 0.05 (**Supplementary Figure S12A**). In many cases, Whippet nodes were associated with undefined PSI values. We considered the corresponding s-exons as unsupported by RNA-Seq data. The PSI values for the canonical and alternative paths defining the ASEs detected in the curated set and documented in the literature were retrieved from Whippet output using the map we defined between ThorAxe s-exons and Whippet nodes. We considered that an ASE was tissue-regulated when the PSI difference between the canonical and alternative paths was higher than 0.15 in some tissue in at least two species (**Supplementary Figure S12B**). If only one of the two paths was present in a tissue, we asked for its PSI value to be above 0.55 (**Supplementary Figure S12C**). The tissue ontology is described in Supplementary Table S8.

### 3D structural analysis

The 3D structural templates were searched and aligned using our new iterative and s-exon-centred version of PhyloSofS molecular modeling routine (Ait-hamlat et al., 2020). They were visualised using Pymol (DeLano, 2002).

### Gene Ontology analysis

The Gene Ontology (GO) annotations for the human proteome were downloaded from the GO Consortium online ressource (Ashburner et al., 2000; Consortium, 2019). We focused on the subset of labels of type “cellular components”. For each label and each gene class, MEX, ALT, REL, UNREL or NO (comprised of the protein-coding genes not included in the other classes), we computed a p-value using a two-sided hypergeometric test (Rivals et al., 2007). We considered that a label was significantly enriched or depleted in a gene class if the p-value was lower than 1*e*^−5^. Generic or vague labels such as “cell” and those containing “organelle” and “part” were excluded from the analysis.

### Comparison with Phastcons scores

We downloaded the human genome Conservation annotation track from the UCSC Genome Browser (https://genome.ucsc.edu). It gives the Phastcons score of each base pair in the human genome computed by the PHAST package (Siepel and Haussler, 2005; Siepel et al., 2005) from the multiple alignments with 99 vertebrate genomes. PhastCons is a hidden Markov model-based method estimating the probability of each nucleotide to belong to a conserved element. It considers not only the column corresponding to the nucleotide of interest in the alignment but also its flanking columns. The PhastCons scores range between 0 and 1. We converted them into residue-based scores by taking the maximum value computed over the 3 nucleotides encoding each residue (<3 for residues overlapping with exon boundaries). Since Phastcons is sensitive to “runs” of conserved sites, we restricted the analysis to s-exons longer than 10 residues. In total, we treated 199,916 s-exons covering 10,060,639 residue positions.

### Comparison with other methods

We bypassed ThorAxe clustering step and/or refinement step by directly modifying the tool’s command line arguments. To compare ThorAxe with the RBBH method, we performed an all-to-all comparison between species. Specifically, given two species *s*_1_ and *s*_2_, we ran BLAST for each sub-exon defined by ThorAxe in *s*_1_ against the ensemble of sub-exons from *s*_2_, and reciprocally, and we identified the best reciprocal pairs of s-exons across the two species. Defining s-exons from a set of pairwise alignments of sub-exons is a difficult task, and thus we simply compared the sub-exon pairs identified by RBBH with those implied by ThorAxe s-exon definition.

## Data access

All data accessed from public repositories – namely Ensembl (www.ensembl.org, GRCh37 assembly), Bgee (*https://bgee.org*), the UCSC Genome Browser (https://genome.ucsc.edu), and the Protein Data Bank (PDB, www.rcsb.org) – are detailed in the Methods. We declare that all data supporting the findings of this study are available within the paper, via the supplementary webserver http://www.lcqb.upmc.fr/ThorAxe and in the Supplementary Information files.

## Acknowledgments

A grant of the French national research agency (MASSIV project, ANR-17-CE12-0009) provided a salary to D.J.Z. and S.L.. We thank S. Grudinin for insightful comments.

## Notes

https://github.com/PhyloSofS-Team/thoraxe

http://www.lcqb.upmc.fr/ThorAxe

### Competing Interest Statement

The authors have declared no competing interest.

### Summary of Updates

We corrected the order of the authors.

