## Supplementary Information for "Assessing Conservation of Alternative Splicing with Evolutionary Splicing Graphs"

### Supplementary Text

#### Algorithm 1

This algorithm identifies a set $s$ of s-exons as contiguous blocks in an input MSA $msa$, where the letters indicate to which sub-exon each residue belongs to. For instance, if $msa[i, j] = "a"$, then it means that the residue at position $i$ in the MSA and coming from sequence $j$ belongs to the sub-exon "a". Whenever there is a change in sub-exon, the algorithm define a new s-exon.

```
s ← {}
start ← 0
for i = 1 to L − 1 do
    if start = 0 then
        start ← i
    end
    c ← 0
    has2stop ← False
    j ← 1
    while j ≤ n and not has2stop do
        if msa[i, j] ≠ " − " and msa[i + 1, j] ≠ " − " then
            if msa[i, j] ≠ msa[i + 1, j] then
                has2stop ← True
            else
                c ← c + 1
            end
        end
    end
    if has2stop or c = 0 then
        stop ← i
        s ← s||{[start, stop]}
        start ← 0
    end
end
if start = 0 then
    start ← L
end
s ← s||{[start, L]}
```

where $L$ and $n$ are the numbers of positions and sequences, respectively, in the MSA.

#### Algorithm 2

This algorithm detects AS events as variations displayed by a set of input transcripts with respect to a reference canonical transcript $c$.

```
forEach input transcript t do
    i ← 2
    j ← 2
    while i ≤ ne(t) do
        if v_t^i ≠ v_c^j then
```

find $k$ and $l$ such that $v_t^k = v_c^l$
$event \leftarrow ([v_c^{(j-1)} : v_c^l],[v_t^{(i-1)} : v_t^k])$
add *event* to the list of detected AS events
$i \leftarrow k + 1$
$j \leftarrow l + 1$
**end**
**end**
**end**

where $ne(t)$ is the number of s-exons in $t$ (or nodes in the path defining $t$), and $v_t^i$ is the *ith* s-exon of $t$ (or the *ith* node in the path defining $t$). Note that the first and last nodes in each transcript path are the *start* and the *stop*.

#### Computational details

ThorAxe v0.6.3 was run on every human protein-coding gene using the following command:

```
thoraxe -i $protein -o $protein -y --plot_chimerics -l $sp
```

where the variable *protein* stores the name of the query gene and *sp* stores the list of species considered. The calculation over the whole proteome completed in 240 hours single-core and about 19 hours using Julia to parallelize the dataset in 15 cores and with the WSL2 of Windows 10, on an Intel(R) Xeon(R) W-2145 CPU @ 3.70GHz. On average, each gene was treated in 56 seconds.
The command used to run ThorAxe without the clustering step was:

```
thoraxe -i $protein -o $protein --coverage 0.0 --identity 0.0 -y --plot_chimerics -l $sp
```

The command used to bypass both the clustering step and the refinement step was:

```
thoraxe -i $protein -o $protein --coverage 0.0 --identity 0.0 -y --plot_chimerics -l $sp \
 --no_movements --no\_disintegration
```

Table S1: **Curated set**

| Main gene | Auxiliary genes | #(species) | Function |
|---|---|---|---|
| BCL2L1 | - | 7 | regulates outer mitochondrial membrane channel opening, and hence apoptosis |
| CAMK2B | CAMK2A,D,G | 4-10 | plays multiple unique roles in actin assembly (organisation, stabilisation, polymerization...) |
| DNM2 | DNM1,3 | 8-10 | GTPase involved in membrane remodelling, engaged in many protein-protein interactions |
| FMR1 | FXR1,2 | 9-11 | multifunctional polyribosome-associated RNA-binding protein |
| FYN | FGR, SRC, YES | 10-11 | tyrosine kinase involved in T-cell and neuronal signaling |
| GRIN1 | GRIN2A,2B,2C,2D,3A,3B | 6-11 | glutamate and ion channel protein receptor activated when glycine and glutamate bind to it |
| KIF1B | KIF1A,C | 10 | microtubule-dependent motor protein involved in cellular trafficking |
| MAPK8 | MAPK9,10 | 9-11 | serine/threonine kinase involved in many essential signaling pathways |
| MYH11 | MYL6,9,12B | 6-10 | major contractile protein involved in muscle contraction |
| MYO1B | MYO1C,D,E,F,G,H | 7-12 | links lipid membrane to the actin cytoskeleton, plays roles in membrane trafficking and dynamics |
| NEBL | - | 8 | Binds to actin and plays an important role in the assembly of the Z-disk |
| Nxnl2 | Nxnl1 | 9-10 | may be involved in the viability of sensory neurons |
| PAX6 | - | 10 | key transcription factor involved in eye development |
| PTPRC | - | 9 | protein tyrosine phosphatase receptor regulating lymphocytes signalling |
| SNAP25 | SNAP23 | 10-11 | Part of the SNARE complex, which is involved in vesicle fusion |
| TPM1 | TPM2,3,4 | 5-7 | actin-binding protein, involved in muscle contraction and cytoskeleton formation |

For each family, we defined a *main* gene and focused on retrieving information from the literature for that gene. The third column indicates the number of species range where one-to-one orthologs were found, for each family. The function descriptions were taken from Uniprot (`https://www.uniprot.org`) and Wikipedia (`https://www.wikipedia.org`).

Table S2: **Documented AS events from the curated set**

| | Gene | Event | Function |
|---|---|---|---|
| 1 | BCL2L1 | 63-aa deletion | function reversion[1] & partner loss[2] |
| 2 | CAMK2A | 11-aa insertion | cellular localisation[3] |
| 3 | CAMK2B | insertion of 43-aa proline-rich repeats | partner acquisition (?)[4] |
| 4 | | 25-aa deletion | protein partner loss[5] |
| 5 | DNM2 | 46-aa mutually exclusive homologous exons | partner specificity[6] |
| 6 | | 4-aa deletion | cellular localisation[6] |
| 7 | FMR1 | 21-aa deletion (ex. 12) | RNA-binding affinity[7] |
| 8 | | 17-aa deletion (ex. 14) | cellular localisation & RNA binding[7] |
| 9 | | 13-aa deletion (alt. acc. in ex. 15) | PTM sites & RNA-binding affinity[7] |
| 10 | | 25-aa deletion (alt. acc. in ex. 15) | PTM sites & RNA-binding affinity[7] |
| 11 | | 17-aa deletion (alt. acc in ex. 17) | cellular localisation[7] |
| 12 | FYN | 50-aa mutually exclusive homologous exons | domain organisation (linker)[8] |
| 13 | GRIN1 | 21-aa deletion (ex. 5) | ligand binding affinity & regulation[9;10] |
| 14 | | alternative end | ligand binding affinity & regulation[9;10] |
| 15 | | 37-aa deletion (ex. 21) | ligand binding affinity & regulation[9;10] |
| 16 | KIF1B | 490 aas (1 ex.) replaced by 1100 aas (27 ex.) | partner binding specificity[11] |
| 17 | | 83-aa deletion | partner binding specificity[11] |
| 18 | | 6-aa deletion in the N-domain | binding affinity & dimerisation[11] |
| 19 | | 40-aa deletion in the N-domain | binding affinity & dimerisation[11] |
| 20 | MAPK8 | 10-aa mutually exclusive homologous exons | substrate selectivity[12] |
| 21 | MYH11 | 7-aa insertion | function regulation[13] |
| 22 | MYO1B | 25-aa insertions and variations | protein binding affinity[14] |
| 23 | NEBL | 12 nebulin repeats replaced by a LIM domain | partner selectivity[15] |
| 24 | Nxnl2 | thioredoxin domain present/absent | novel function in signaling[16] |
| 25 | PAX6 | 14-aa insertion | DNA-binding specificity[17] |
| 26 | | paired-domain deletion | DNA-binding specificity[17] |
| 27 | PTPRC | up to 3 exon-deletions | immunological recognition[18] |
| 28 | SNAP25 | 34-aa mutually exclusive homologous exons | partner specificity (?)[19] |
| 29 | TPM1 | 26-aa mutually exclusive homologous exons | partner affinity and dissociation[20] |
| 30 | | 3 ex. deletion coupled with replacement | partner affinity and dissociation[20] |

The symbol "?" in the last column indicates that the functional annotation of the ASE remains speculative. The row colors indicate the level of evidence from gene annotations and RNA-Seq splice junctions. Green: supported by both the annotations and the RNA-Seq data, and tissue-regulated. Light green: supported by both the annotations and the RNA-Seq data, without any evidence of tissue regulation. Light blue: detected only in the annotations. Grey: not detected. See Supplementary Table S3 for more detailed information.

Table S3: **Detection of the documented AS events by ThorAxe**

| | **Gene** | **Number of species** | **Event rank** | **Number of s-exons** | $SF$ | | $ATF$ | | $SF^{RNASeq}$ | | tissue regulation |
|---|---|---|---|---|---|---|---|---|---|---|---|
| | | | | | *can* | *alt* | *can* | *alt* | *can* | *alt* | |
| 1 | BCL2L1 | 7 | 1 | 2 | 100 | 29 | 69 | 12 | 29 | 29 | |
| 2 | CAMK2A | 10 | 1 | 1 | 100 | 70 | 51 | 28 | 70 | 70 | ✓ |
| 3 | CAMK2B | 10 | 2 | 4 | 100 | 70 | 46 | 22 | 80.0 | 50.0 | ✓ |
| 4 | | | 3 | 2 | 50 | 40 | 25 | 14 | 50 | 40 | ✓ |
| 5 | DNM2 | 10 | 2 | 2/2 | 100 | 60 | 77 | 17 | 100 | 60 | ✓ |
| 6 | | | 1 | 1 | 100 | 80 | 57 | 32 | 100 | 100 | ✓ |
| 7 | FMR1 | 11 | 1 | 1 | 82 | 64 | 42 | 24 | 73 | 73 | ✓ |
| 8 | | | - | - | - | - | - | - | - | - | |
| 9 | | | - | - | - | - | - | - | - | - | |
| 10 | | | 3 | 2 | 91 | 27 | 66 | 8 | 82 | 45 | ✓ |
| 11 | | | 5 | 1 | 73 | 45 | 45 | 30 | 36 | 36 | ✓ |
| 12 | FYN | 11 | 1 | 1/1 | 100 | 73 | 52 | 32 | 91 | 73 | ✓ |
| 13 | GRIN1 | 10 | 1 | 1 | 90 | 70 | 62 | 28 | 90 | 90 | ✓ |
| 14 | | | 2 | 2/1 | 60 | 60 | 27 | 21 | 20 | 0 | |
| 15 | | | 3 | 3 | 50 | 40 | 23 | 15 | 10 | 0 | |
| 16 | KIF1B | 10 | 5 | 37/1 | 40 | 60 | 15 | 18 | 0 | 30 | |
| 17 | | | 3 | 1 | 100 | 40 | 61 | 9 | 80 | 80 | ✓ |
| 18 | | | 1 | 1 | 80 | 70 | 51 | 32 | 70 | 70 | ✓ |
| 19 | | | 4 | 2 | 40 | 40 | 17 | 14 | 40 | 40 | ✓ |
| 20 | MAPK8 | 9 | 1 | 1/2 | 89 | 89 | 38 | 36 | 89 | 78 | ✓ |
| 21 | MYH11 | 10 | 1 | 1 | 90 | 90 | 44 | 40 | 80 | 70 | ✓ |
| 22 | MYO1B | 9 | 1 | 2 | 89 | 89 | 38 | 23 | 78 | 89 | ✓ |
| 23 | NEBL | 8 | 2 | 24/4 | 63 | 50 | 19 | 13 | 0 | 0 | |
| 24 | Nxnl2 | 10 | 1 | 1/1 | 89 | 22 | 80 | 15 | 10 | 0 | |
| 25 | PAX6 | 10 | 1 | 1 | 70 | 70 | 25 | 31 | 50 | 50 | ✓ |
| 26 | | | 4 | 4 | 30 | 20 | 12 | 13 | 0 | 0 | |
| 27 | PTPRC | 9 | 1 | 3 | 33 | 33 | 11 | 7 | 11 | 22 | ✓ |
| 28 | SNAP25 | 11 | 1 | 1/1 | 82 | 64 | 48 | 34 | 82.0 | 64.0 | ✓ |
| 29 | TPM1 | 7 | 1 | 1/1 | 100 | 100 | 52 | 40 | 86 | 71 | ✓ |
| 30 | | | 2 | 3/1 | 86 | 57 | 45 | 26 | 0 | 0 | |

The rank of an event reflects its conservation level with respect to the other events detected for the gene. $SF$ and $ATF$ are the species and averaged transcript fractions (given in percentages) computed by ThorAxe for the canonical (*can*) and alternative (*alt*) paths. $SF^{RNASeq}$ gives the proportion of species where the presence of the canonical or alternative subpath is supported by RNA-Seq data. Tissue regulation must be observed for at least one tissue in at least two species to be considered.

Table S4: **Per-gene statistics computed over all human protein coding genes**

| **Variable** | **Mean** | **Std** | **Min** | **q25** | **Median** | **q75** | **Max** |
|---|---|---|---|---|---|---|---|
| Species | 8.5 | 2.6 | 1 | 7 | 9 | 10 | 12 |
| Transcripts | 17.1 | 10.3 | 1 | 10 | 15 | 22 | 113 |
| Exons | 103.1 | 102.2 | 1 | 31 | 74 | 142 | 1677 |
| Sub-exons | 102.3 | 102.2 | 1 | 31 | 73 | 140 | 1685 |
| S-exons | 25.8 | 25.2 | 1 | 9 | 18 | 35 | 354 |

Table S5: **List of MEX and ALT s-exons.** See the supplementary data *datPairs_MEX_ALT.csv*. The s-exons are grouped by ASE.

Table S6: **List of REL s-exons.** See the supplementary data *datPairs_REL.csv*. The s-exons are grouped by ASE.

Table S7: **List of UNREL s-exons.** See the supplementary data *datPairs_UNREL.csv*. The s-exons are grouped by ASE.

Table S8: **Tissue ontology**

| **Tissue Group** | **Tissue** |
|---|---|
| Ammon's horn | Ammon's horn |
| immune | CD4-positive helper T cell<br>leukocyte<br>spleen |
| immune/thyroid | head kidney<br>lymph node<br>thymus |
| kidney | adult mammalian kidney |
| bone | bone tissue |
| brain | brain<br>cerebellum<br>frontal cortex<br>head<br>prefrontal cortex |
| digestive | colon<br>intestine<br>liver<br>stomach |
| embryo | blastula<br>embryo<br>gastrula<br>placenta |
| eye | eye |
| female reproduction | female gonad<br>mature ovarian follicle |
| heart | heart<br>heart left ventricle |
| kidney | kidney<br>mesonephros |
| lung | lung |
| male reproduction | prostate gland<br>testis |
| multi-cellular organism | multi-cellular organism |
| muscle | muscle of leg<br>muscle tissue<br>skeletal muscle tissue |
| pharyngeal gill | pharyngeal gill |
| thyroid gland | thyroid gland |
| zone of skin | zone of skin |

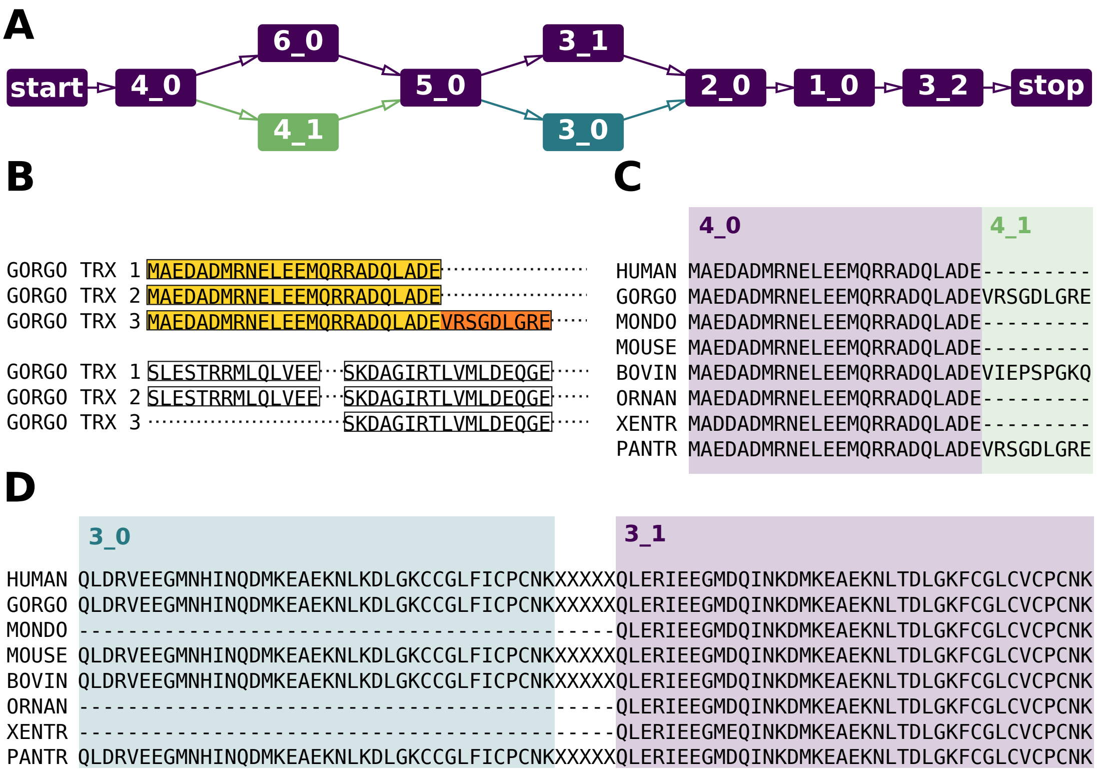


Figure S1: **Examples of ThorAxe intermediate and final outputs. A**. Evolutionary splicing graph computed for SNAP-25 across 8 species. Each node represents a s-exon, and two nodes are linked by an edge if they are consecutive in at least one input transcript. The edges and the nodes are colored according to their conservation level (species fraction, see *Materials and Methods*). For ease of visualisation, the species-specific s-exons (*i.e.* defined in only one species) were filtered out. **B**. Redundancy reduction step illustrated with three transcripts observed in gorilla. From two overlapping exons, one shorter and one longer, we define two sub-exons, highlighted in yellow and orange. **C-D**. MSAs associated to the exon clusters numbered 3 and 4. The final output s-exons (*3_0*, *3_1*, *4_0*, *4_1*) are detected as contiguous blocks in the MSAs. The colors of the blocks match those of the nodes in panel A. The s-exons 3_0 and 3_1 are mutually exclusive and their sequences are highly similar.

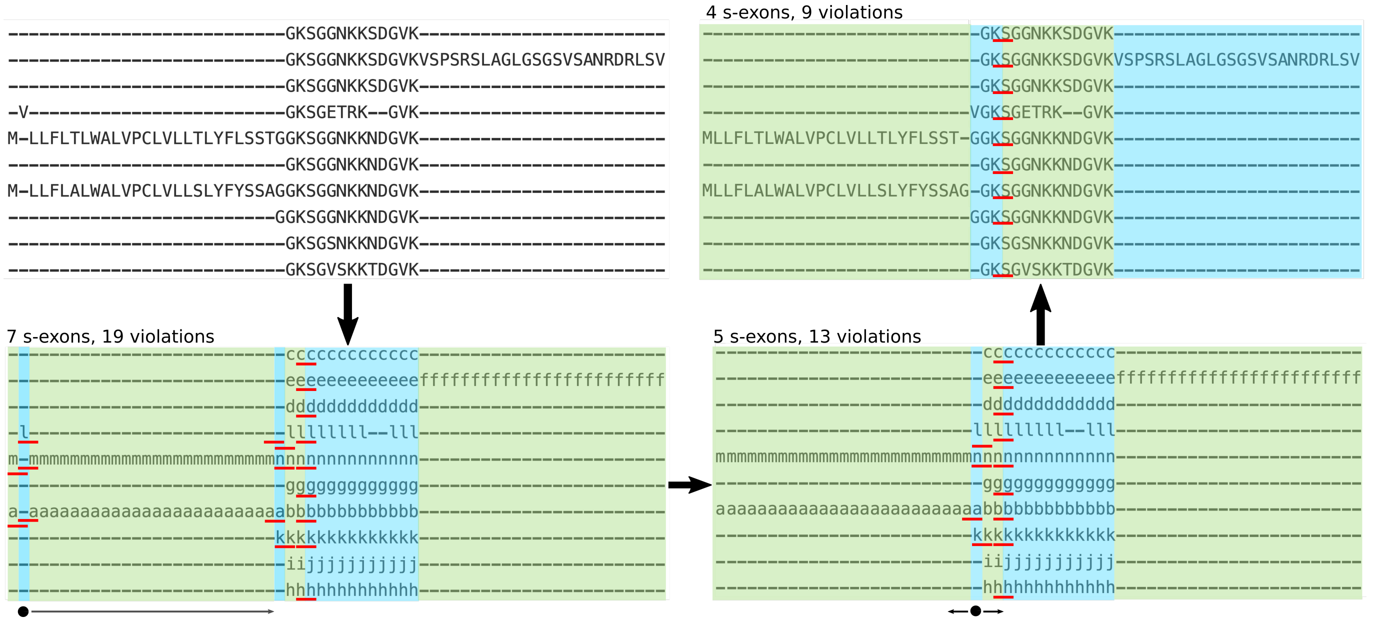


Figure S2: **Refinement procedure.** The top left panel shows a chimeric MSA, where each letter corresponds to an amino acid residue. The bottom left panel shows the translation of this MSA into the sub-exon alphabet. In other words, each letter indicates to which sub-exon the residue at this position belongs to. The coloured blocks correspond to the s-exons defined from the MSA. There are 7 s-exons, among which 3 very small ones (1- or 2-column long) arising from inconsistencies between sub-exon boundaries. This is highlighted by the red lines, which indicate the s-exon boundaries splitting the sub-exons. We refer to those cases as *violations*, and there are 19 of them in this MSA. ThorAxe refinement procedure locally optimises the MSA by first moving the 1-residue long stretch "V" (indicated by a black dot) to the right in order to group it with the rest of its sub-exon, denoted "l". The bottom right panel shows that this operation reduces the number of s-exons to 5 and the number of violations to 13. Second, the remaining 1-residue long s-exon (indicated by a black dot) is disintegrated by separating sub-exons "l", "n" and "k" from sub-exon "a". In practice, the "m"- and the "a"- stretches are shifted to the left in the MSA. This leads to the solution displayed in the top right panel, with only 4 s-exons and 9 violations.

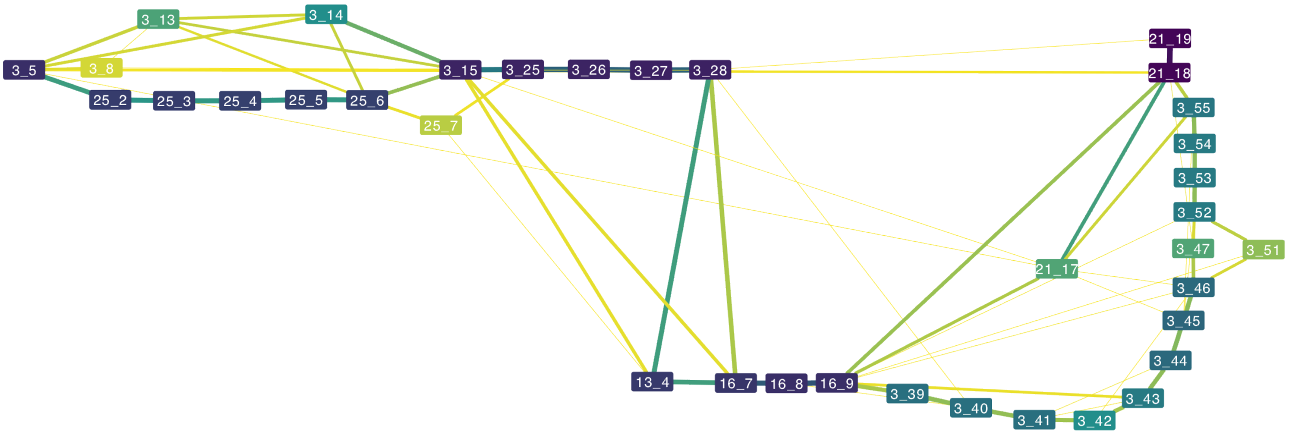


Figure S3: **CAMK2B linker transcript variability across a very large number of species.** The ESG was computed by ThorAxe starting from 499 transcripts annotated in 93 species. These are all the species annotated in Ensembl where one-to-one orthologs could be found. The s-exons present in less than 10% of the species were removed to ease visualisation.

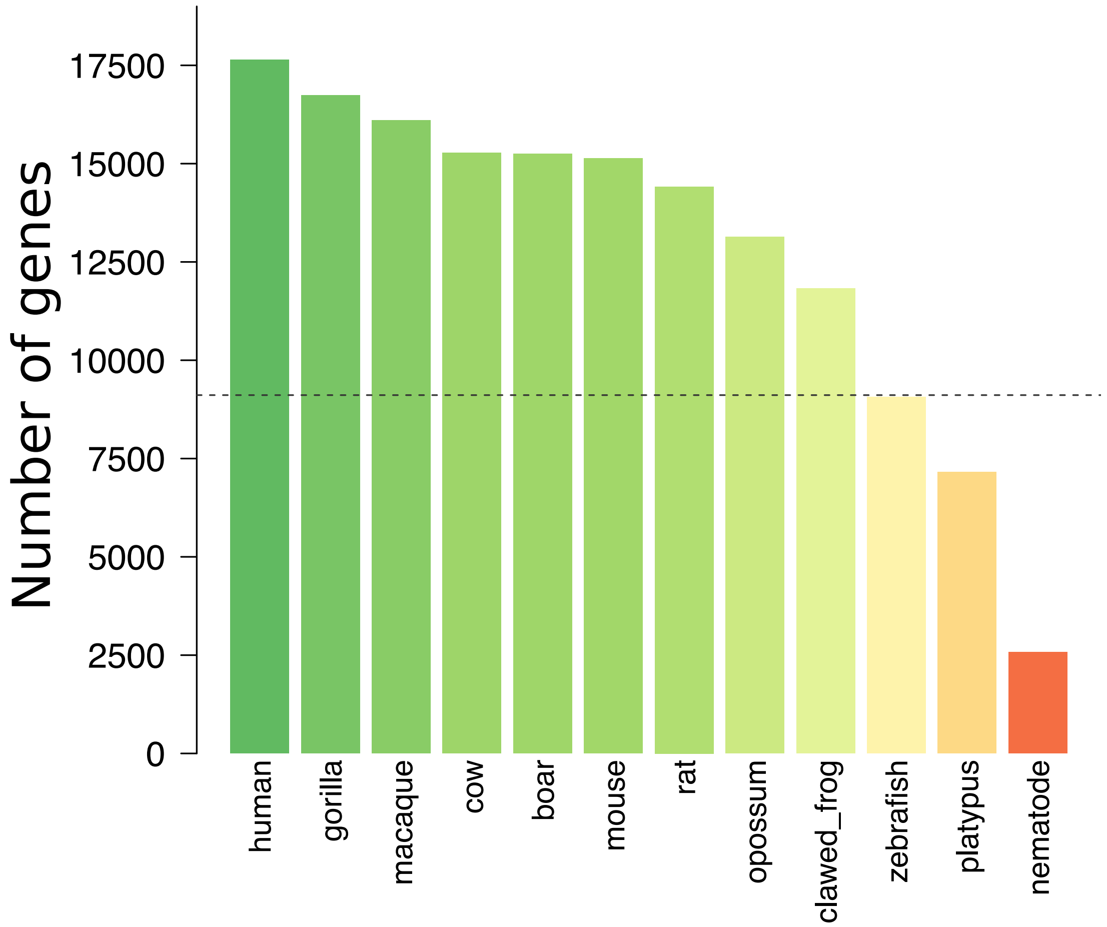


Figure S4: **Number of genes within each considered species.** The color code goes from green (many genes) through yellow to red (few genes).

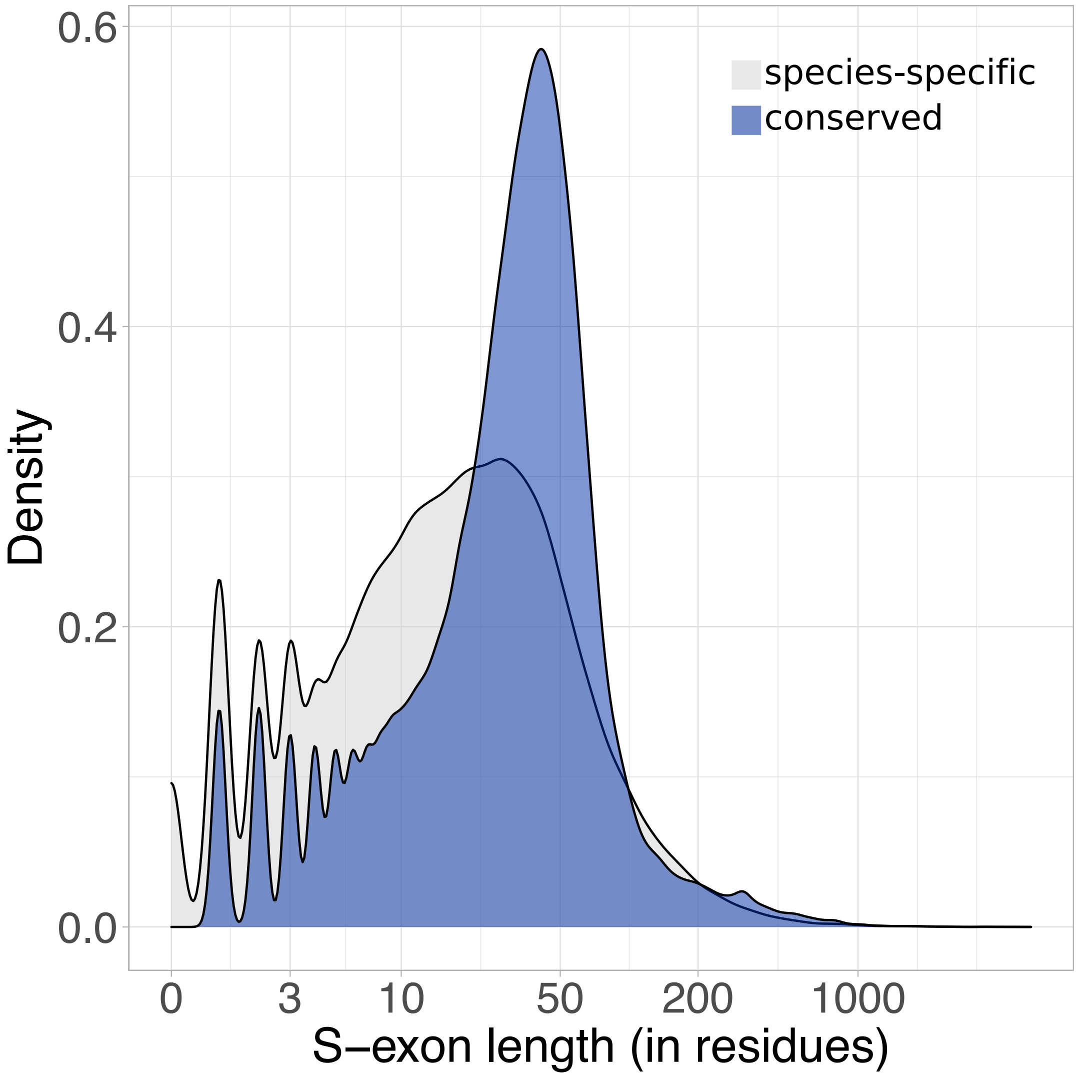


Figure S5: **S-exon length.** Distribution densities of the s-exon lengths. The length of a s-exon is the maximum number of amino acid residues determined over the sequences comprised in the MSA (see *Materials and Methods*). The s-exons are classified as either species-specific (only one sequence in the MSA) or conserved (more than one sequence).

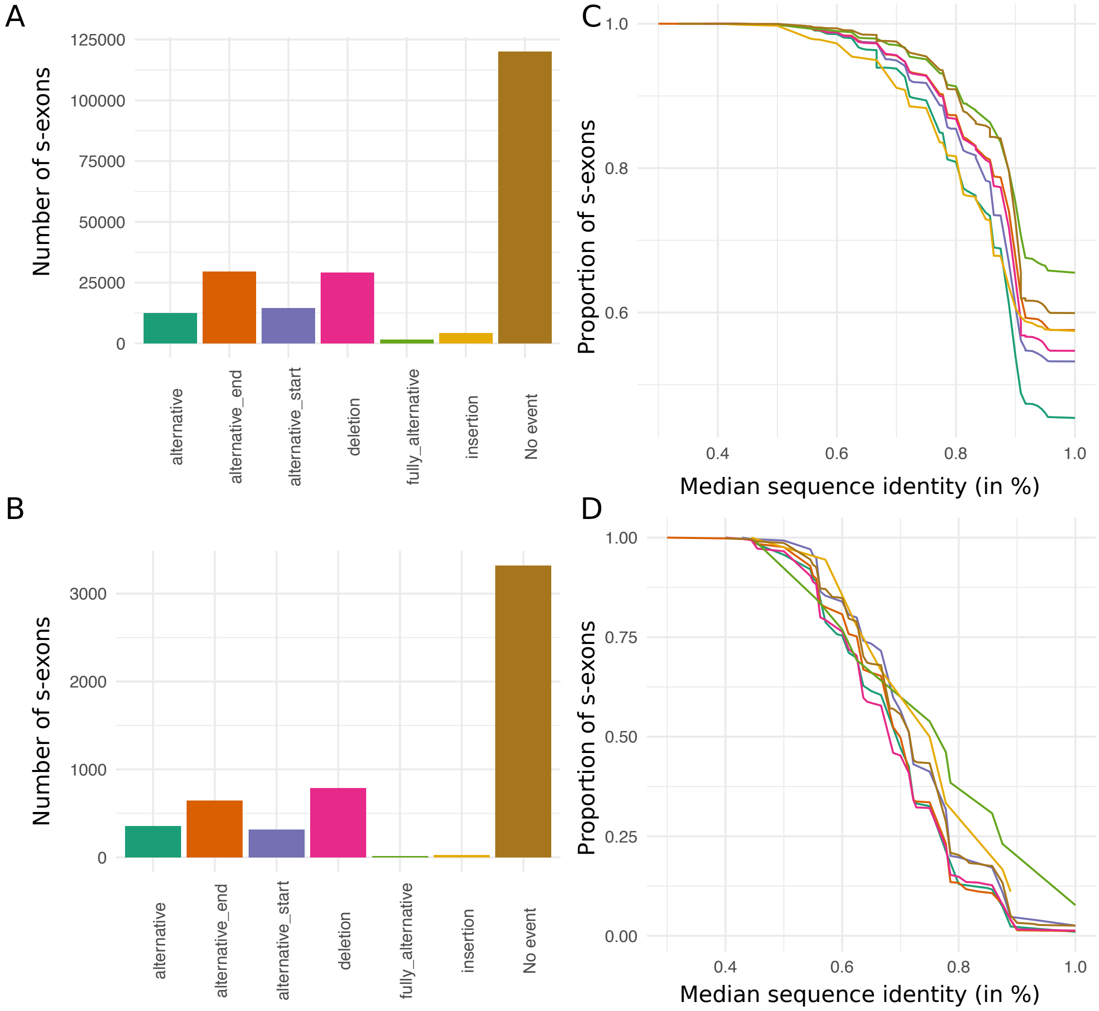


Figure S6: **S-exon involvement in ASEs and Phastcons scores. (A-B)** Barplots of the numbers of s-exons involved in the different types of ASEs, and those not involved in any ASE. **(C-D)** Cumulative distributions of MSA sequence identity percentage. On the y-axis we report the percentage of s-exons with a median column identity greater than the x-axis value. **(A,C)** All s-exons longer than 10 residues and belonging to genes with one-to-one orthologs in at least eight species. **(B,D)** Selection of s-exons displaying high species fractions (>0.8) but low Phastcons median scores (<0.1).

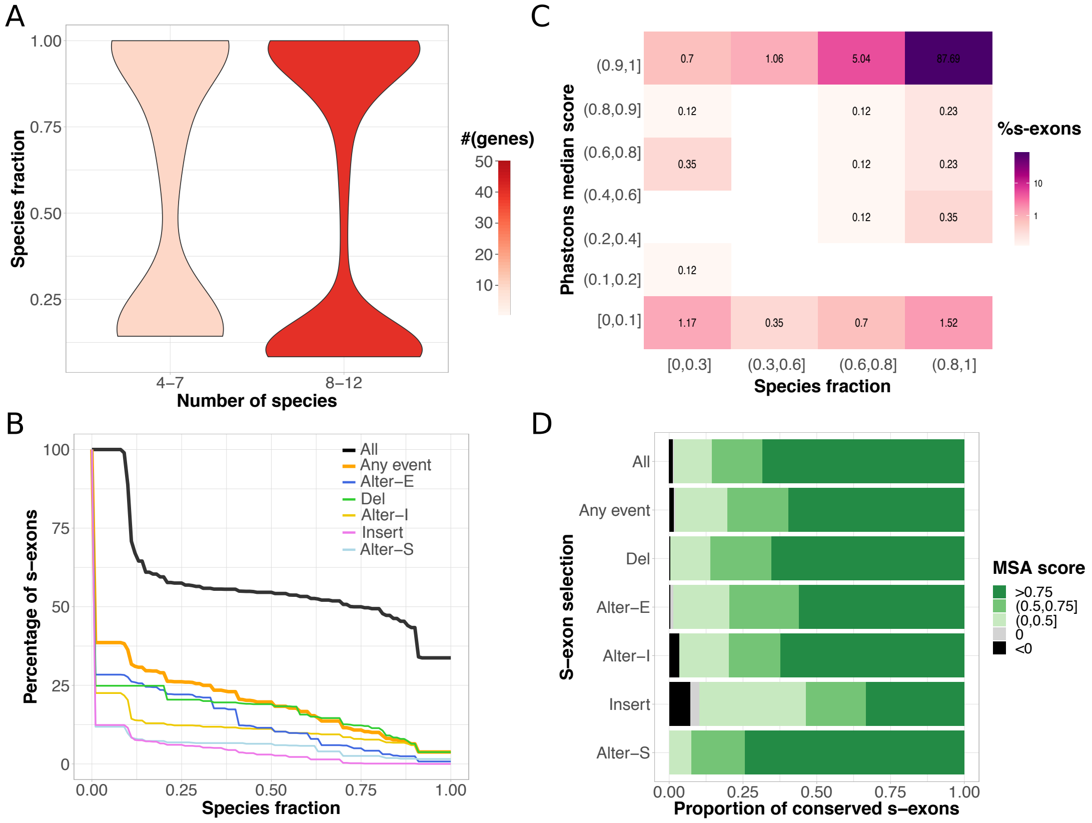


Figure S7: **Properties of the s-exons and AS events detected over our curated set of 50 genes.** **A**. Distributions of the s-exon species fractions depending on the number of species. The red tones indicate the number of genes comprised in each distribution. **B**. Cumulative distributions of s-exon species fraction. On the y-axis we report the percentage of s-exons with a species fraction greater than the x-axis value. The different curves correspond to all s-exons (*All*), only those involved in at least an ASE (*Any event*), or only those involved in a specific type of event. *Alter-S*: alternative start. *Alter-I*: alternative (internal). *Alter-E*: alternative end. *Del*: deletion. *Insert*: insertion. **C**. Heatmap of the s-exon Phastcons median scores versus the s-exon species fractions. Only the s-exons longer than 10 residues and belonging to genes with one-to-one othologs in at least 8 species are shown. **D**. Proportions of conserved s-exons displaying very poor (negative score) to very good (score close to one) alignment quality. The MSA score of a s-exon is computed as a normalised sum of pairs. A score of 1 indicates 100% sequence identity without any gap. The proportions are given for different s-exon selections (same labels as in panel B).

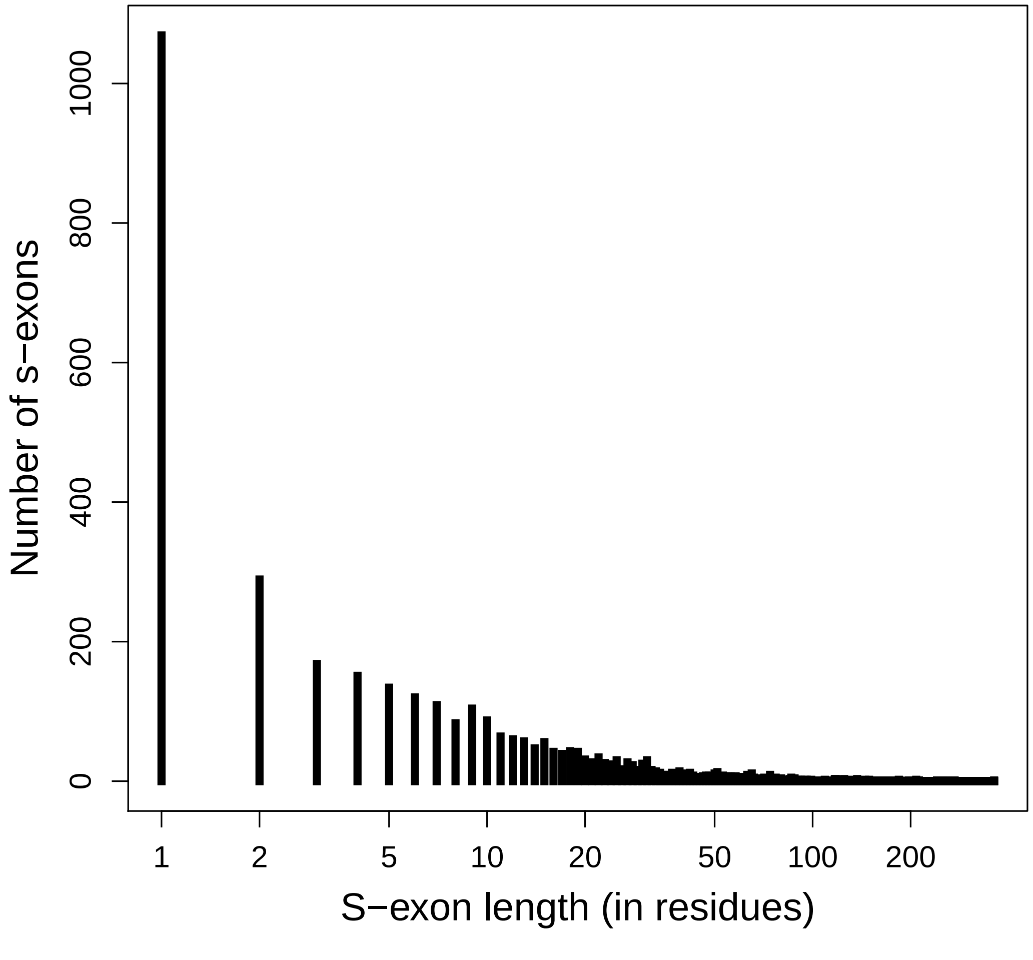


Figure S8: **Length distribution for the s-exons defined by very poor quality MSAs (negative score).**

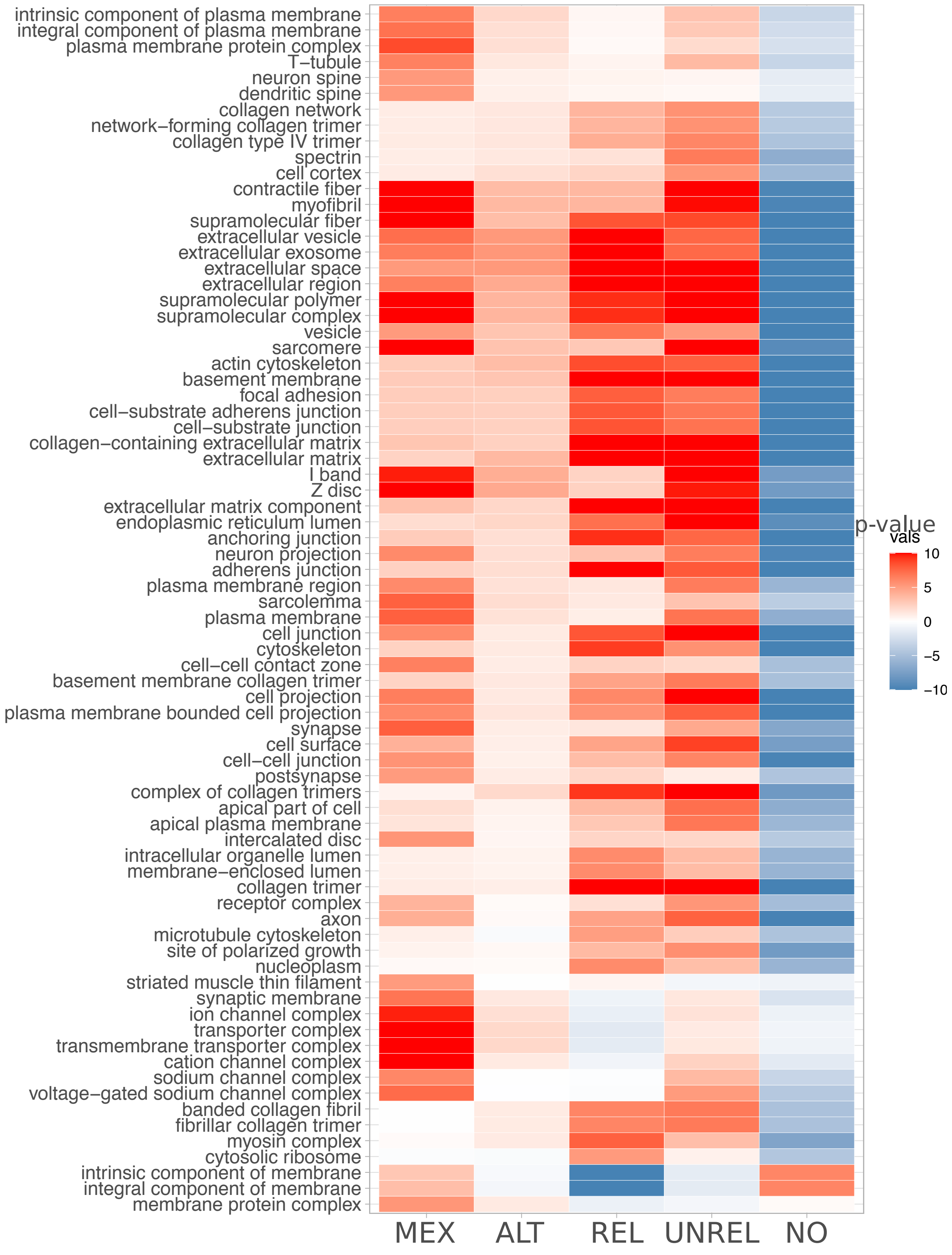


Figure S9: **Cellular localisation of the genes where similar sequences are alternatively used.** The genes are classified according to the role of the detected similar s-exons in ASEs. MEX: mutually exclusive s-exons. ALT: alternative (non mutually exclusive) s-exons. REL: one s-exon is in the canonical or alternative subpath of an event (of any type), while the other one serves as a "canonical anchor" for the event. UNREL: one s-exon is in the canonical or alternative subpath of an event (of any type), while the other one is located outside the event in the canonical transcript. NO: no s-exon pair detected. We report the set of Gene Ontology labels of type "cellular component" that are significantly enriched in at least one of the considered gene classes.

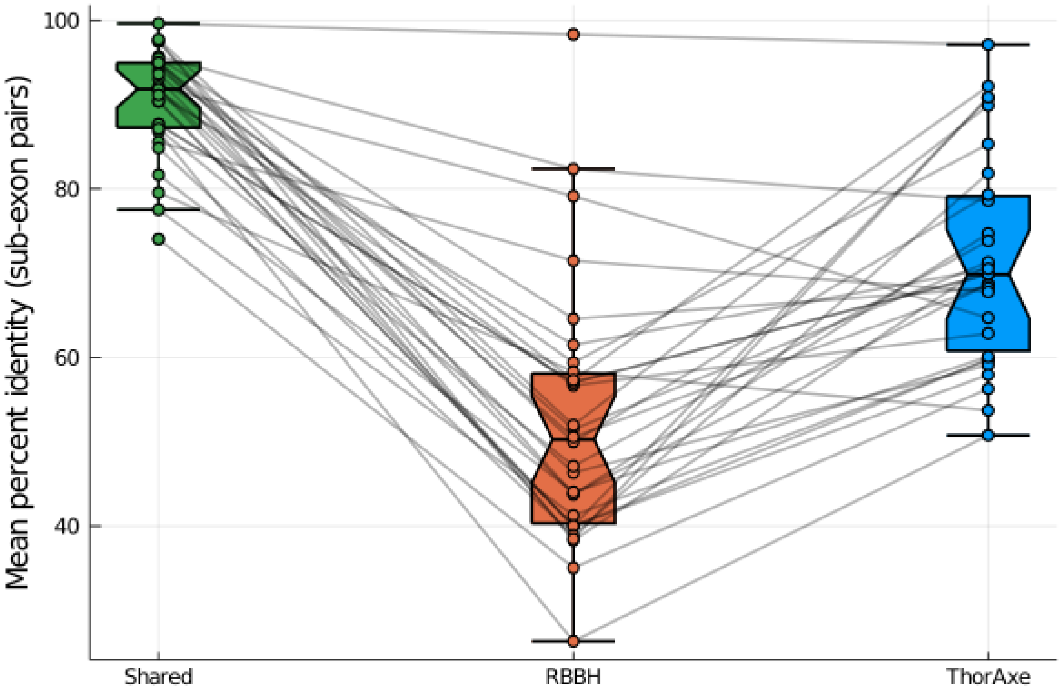


Figure S10: **Sequence identity of the sub-exon pairs detected by ThorAxe and RBBH.** The identity percentages are computed for all the sub-exon pairs detected in the curated set and averaged by gene. 86.98% of the pairs are shared (detected by both approaches), 11.51% are discovered only by ThorAxe, and only 1.51% are RBBH specific.

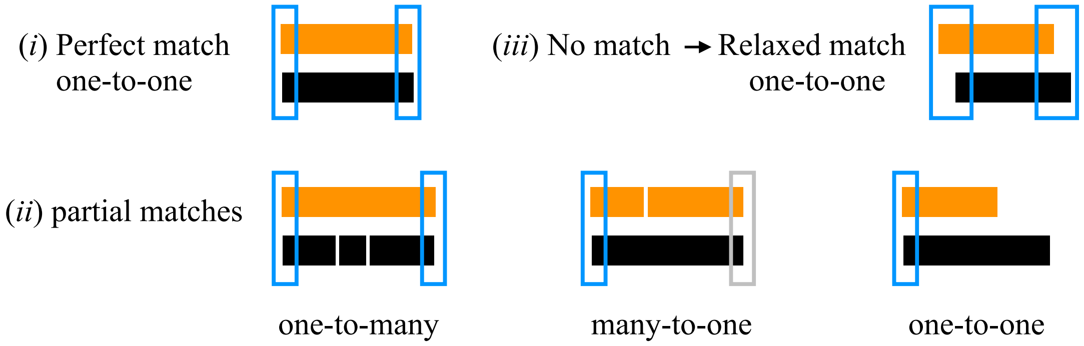


Figure S11: **Types of mapping between ThorAxe s-exons and Whippet nodes.** The orange rectangles represent species-specific sequences extracted from s-exons. In each panel, the query s-exon is the first one. The black rectangles represent Whippet matching nodes sequences. The empty rectangles indicate matches, in blue for the query s-exon and in grey for some other s-exon.

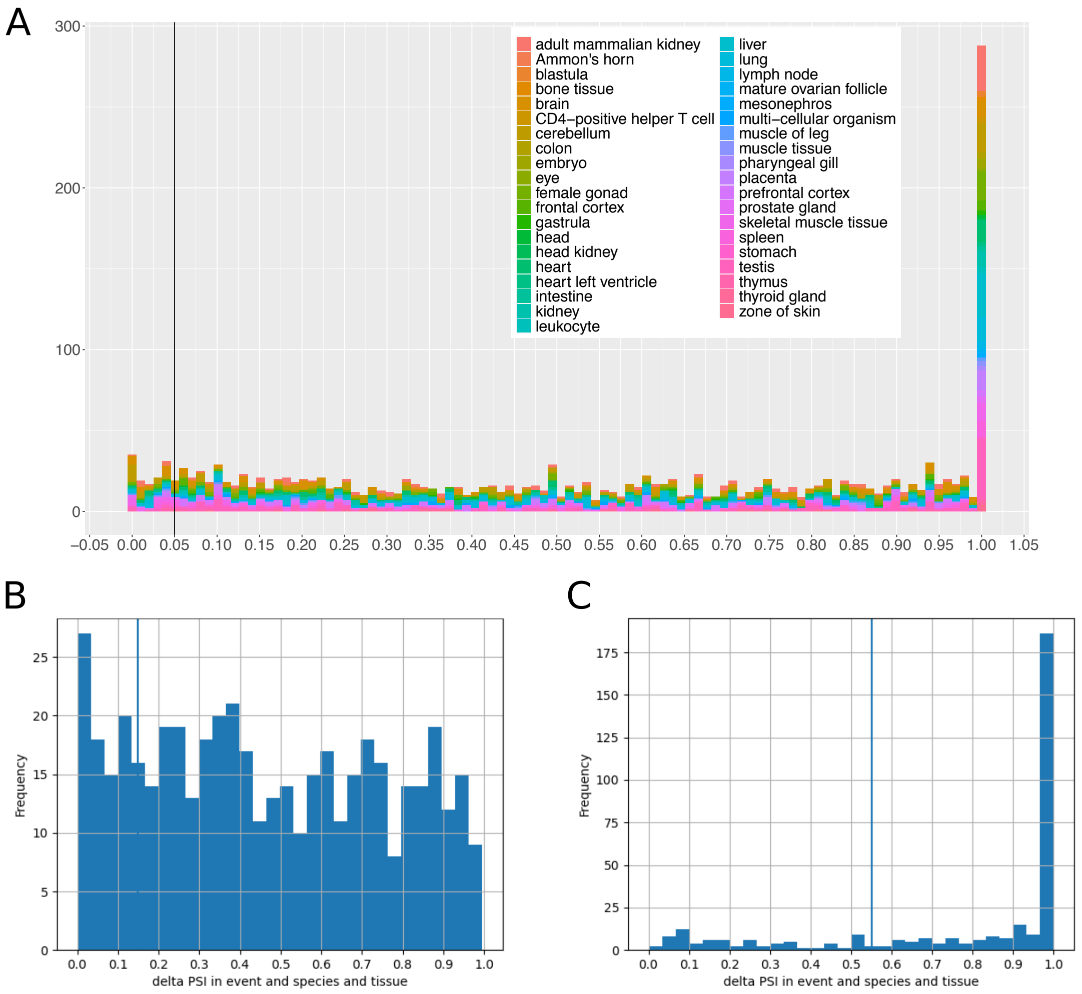


Figure S12: **RNA-Seq Percent Spliced In statistics.** **A**. Distribution of the PSI values computed for the canonical and/or alternative subpaths defining the ASEs detected in the curated set and documented in the literature. The colors indicate the tissues. **B**. Distribution of PSI absolute differences between the canonical and alternative paths, when both are present in the same tissue and species. **C**. Distribution of PSI values for either the canonical or the alternative path, when only one of the two is present in a tissue from a species.
